# Active maintenance of CD8^+^ T cell naïvety through regulation of global genome architecture

**DOI:** 10.1101/2023.02.26.530139

**Authors:** Brendan E. Russ, Kirril Tsyganov, Sara Quon, Bingfei Yu, Jasmine Li, Jason K. C. Lee, Moshe Olshansky, Zhaohren He, Paul F. Harrison, Adele Barugahare, Michael See, Simone Nussing, Alison E. Morey, Vibha A. Udupa, Taylah .J Bennett, Axel Kallies, Cornelis Murre, Phillipe Collas, David Powell, Ananda W. Goldrath, Stephen J. Turner

## Abstract

The differentiation of naïve CD8^+^ cytotoxic T lymphocytes (CTLs) into effector and memory states results in large scale changes in transcriptional and phenotypic profiles. Little is known about how large-scale changes in genome organisation reflect or underpin these transcriptional programs. We utilised Hi-C to map changes in the spatial organisation of long-range genome contacts within naïve, effector and memory virus-specific CD8^+^ T cells. We observed that the architecture of the naive CD8^+^ T cell genome was distinct from effector and memory genome configurations with extensive changes within discrete functional chromatin domains. However, deletion of the BACH2 or SATB1 transcription factors was sufficient to remodel the naïve chromatin architecture and engage transcriptional programs characteristic of differentiated cells. This suggests that the chromatin architecture within naïve CD8^+^ T cells is preconfigured to undergo autonomous remodelling upon activation, with key transcription factors restraining differentiation by actively enforcing the unique naïve chromatin state.

**One Sentence Summary:** CD8^+^ T cell naïvety is actively maintained by transcription factors that enforce a distinct, naïve chromatin architecture.

**Highlights:**

- CD8^+^ T cell differentiation states are underscored by distinct chromatin looping architectures.
- Chromatin loops juxtapose CTL state appropriate enhancers, transcription factors and genes.
- Effector and memory CTLs have similar genome architectures, explaining rapid memory recall.
- CTL differentiation is restrained by BACH2 and SATB1, which enforce a naïve loop architecture.

## INTRODUCTION

Activation of naïve CD8^+^ T lymphocytes triggers a program of clonal expansion and differentiation, resulting in a large pool of effector cells that are then capable of killing virus infected cells via secretion of cytotoxic molecules (granzymes A, B and K, and perforin) (Jenkins et al., 2007). The generation of effector cytotoxic T lymphocytes (CTLs) coincides with the expression of pro-inflammatory chemokines such as CCL4 (MIP1α), CCL5 (RANTES) (Crawford et al., 2011; Russ et al., 2017), Interferon-γ (IFNG) and Tumor Necrosis Factor (TNF) (La Gruta et al., 2004). Upon resolution of infection, a long-lived pool of virus-specific (memory) CTLs is established that, relative to naïve CTLs, elicit effector functions rapidly following re-infection, thus providing the basis of T cell-mediated immunity to subsequent infection (Kaech et al., 2002; Lalvani et al., 1997; Veiga-Fernandes et al., 2000). While it is well established that the different phenotypes and functional capacities of naïve, effector and memory T cells are underscored by unique transcriptomes (Kaech et al., 2002; Russ et al., 2014), how these transcriptional profiles arise and are maintained is not fully understood.

Within eukaryotic cells, DNA is associated with histone protein complexes (nucleosomes), and this association is termed chromatin (Kouzarides, 2007). Changes to the structure of chromatin result in coordinated changes in gene transcription that underly the processes of cellular differentiation, including lineage commitment and acquisition of lineage identity within developing and mature immune cells (Johanson et al., 2018; Russ et al., 2014; Scott-Browne et al., 2016; Yu et al., 2017; Zhang et al., 2012). These changes include modulation of chromatin composition and accessibility at gene regulatory elements such as gene promoters and transcriptional enhancers. Enhancers act as targets for transcription factor (TF) binding that can then directly and indirectly activate or repress gene transcription (Barski et al., 2007; Russ et al., 2014; Russ et al., 2017; Zhang et al., 2012). For instance, TFs including TBET (Intlekofer et al., 2005), BLIMP1 (Rutishauser et al., 2009), and IRF4 (Man et al., 2013) drive acquisition of effector function within CD8^+^ T cells, while TCF1 (Danilo et al., 2018) and FOXO1 (Delpoux et al., 2021; Kerdiles et al., 2009) are required to maintain the quiescence and stemness of naïve T cells. Importantly, transcriptional networks that drive alternate differentiation states act in opposition to enable maintenance of cellular identity. For instance, FOXO1 contributes to CD8^+^ T cell naïvety by driving expression of BACH2 (Delpoux et al., 2021). BACH2, in-turn, limits effector CTL differentiation by occupying enhancers and promoters of CTL effector lineage determining genes that would otherwise be bound by the AP-1 family of TFs. This effectively inhibits transcriptional activation of genes such as *Prdm1* (which encodes BLIMP1) (Roychoudhuri et al., 2016).

Enhancers can occur kilobases to megabases from the genes that they regulate, conveying their effects on gene transcription via looping of the chromatin fibre to bring enhancers and their target gene promoters into close proximity (Bulger and Groudine, 2011; Heintzman et al., 2007). This interaction likely allows the regulatory modules (TFs and chromatin modifying proteins) assembled at the enhancer to access the promoter. For instance, TBET binds to several enhancers at the *Ifng* locus in CD4^+^ T cells and CTLs, where it drives induction of *Ifng* transcription following activation (Intlekofer et al., 2005). TBET in-turn recruits the histone H3K27 demethylases, KDM6B (Jumonji) and KDM6A (UTX), and the SET7/9 histone methyltransferase complex, which mediates methylation of H3K4. Together these activities result in remodelling of the *Ifng* locus, such that the repressive H3K27Me3 modification is removed, and H3K4me2 and H3K4me3 modifications are deposited across the locus (Li et al., 2021; Miller et al., 2010; Miller and Weinmann, 2010). TBET also recruits CCCTC-binding factor (CTCF), which mediates loop formation, including at the *Ifng* locus (Sekimata et al., 2009). TF dependent chromatin structuring that enables enhancer:promoter interactions has been shown to control acquisition of other lineage-specific genes in T cells, including *Il2* (Li et al., 2017), and *Il4*/*Il5*/*Il13* (Agarwal and Rao, 1998; Avni et al., 2002; Zheng and Flavell, 1997). More recently, it was demonstrated that TCF-1 and LEF-1 are critical for ensuring naïve CD8^+^ T cell identify by maintaining a 3-dimensional genome organisation that represses expression of non-CD8^+^ T cell lineage genes (Shan et al., 2021). Hence, lineage fidelity is maintained at the level of chromatin architecture, and localised chromatin restructuring coincides with T cell differentiation and acquisition of lineage-specific function. However, the extent to which reorganisation of *cis*-regulatory elements modulates CD8^+^ T cell differentiation remains unknown. Here, we aimed to address this question by mapping genome-wide *cis*-regulatory interactions to determine how these underpin functional and phenotypic characteristics during virus-specific CTL differentiation.

## RESULTS

### Stable gross-scale genome architecture is observed at distinct stages of virus specific CTL differentiation

We and others have reported that CD8^+^ T cell differentiation is associated changes in histone biochemical modifications, and chromatin accessibility (Araki et al., 2009; Russ et al., 2014; Russ et al., 2017; Scott-Browne et al., 2016; Sen et al., 2016; Yu et al., 2017). However, these data do not provide information about changes in the spatial organisation of chromatin, particularly those involving non-coding regulatory elements. To determine if acquisition and maintenance of CTL lineage function following virus infection is linked to changes in global chromatin architecture, we performed *in situ* HI-C (Rao et al., 2014), utilising adoptive transfer of naïve (CD44^lo^CD62L^hi^) OT-I TCR transgenic CD8^+^ T cells (CD45.1^+^) specific for the ovalbumin peptide (OVA257-264), followed by intranasal (i.n.) infection with the Influenza A/HKx31-OVA virus (Jenkins et al., 2006). Virus-specific CTLs were isolated at effector (d10) and memory (>d60) time-points p.i. for comparison with naïve OT-1s. Further, data from CD4^+^CD8^+^ (double-positive; DP) thymocytes was captured to enable an ontogenically defined context for comparison of our virus-specific CTL datasets. In total we mapped 2.17 billion contacts across the 4 cell states, corresponding to a total of 55,960 unique chromatin loops (**Table S1**).

We initially assessed gross genome organisation by calculating eigenvectors at 100kb resolution and allocating regions into either A or B genomic compartments, which broadly reflect the spatial separation of active and repressed chromatin regions, respectively (Rao et al., 2014). To validate our compartment assignments, we overlaid ATAC-seq data performed on matching samples, finding that chromatin accessibility was enriched within regions of the genome assigned to the A compartment in naïve, effector and memory CTLs, as expected (**Figures S1A and S1B**). Further, a similar relationship was found for histone modifications that identify enhancers and gene promoters (H3K4me2 and H3K4me3, respectively), while the repressive H3K27me3 modification was more evenly distributed across the A and B compartments (**Figure S1A).** While gross changes in compartmentalisation were not observed between differentiation states (**Figure S1C)**, some small-scale transitions were identified (**Figure 1A**). For example, 290 genes moved from the A compartment to B compartment and 773 genes moved from B to A compartments upon naïve to effector CD8^+^ T cell differentiation. However, these changes in compartmentalisation were not associated with changes in gene transcription (**Figure 1B**). Thus, movement of genes between compartments is not a significant means by which CTL differentiation is regulated following virus infection.

**Figure 1.**
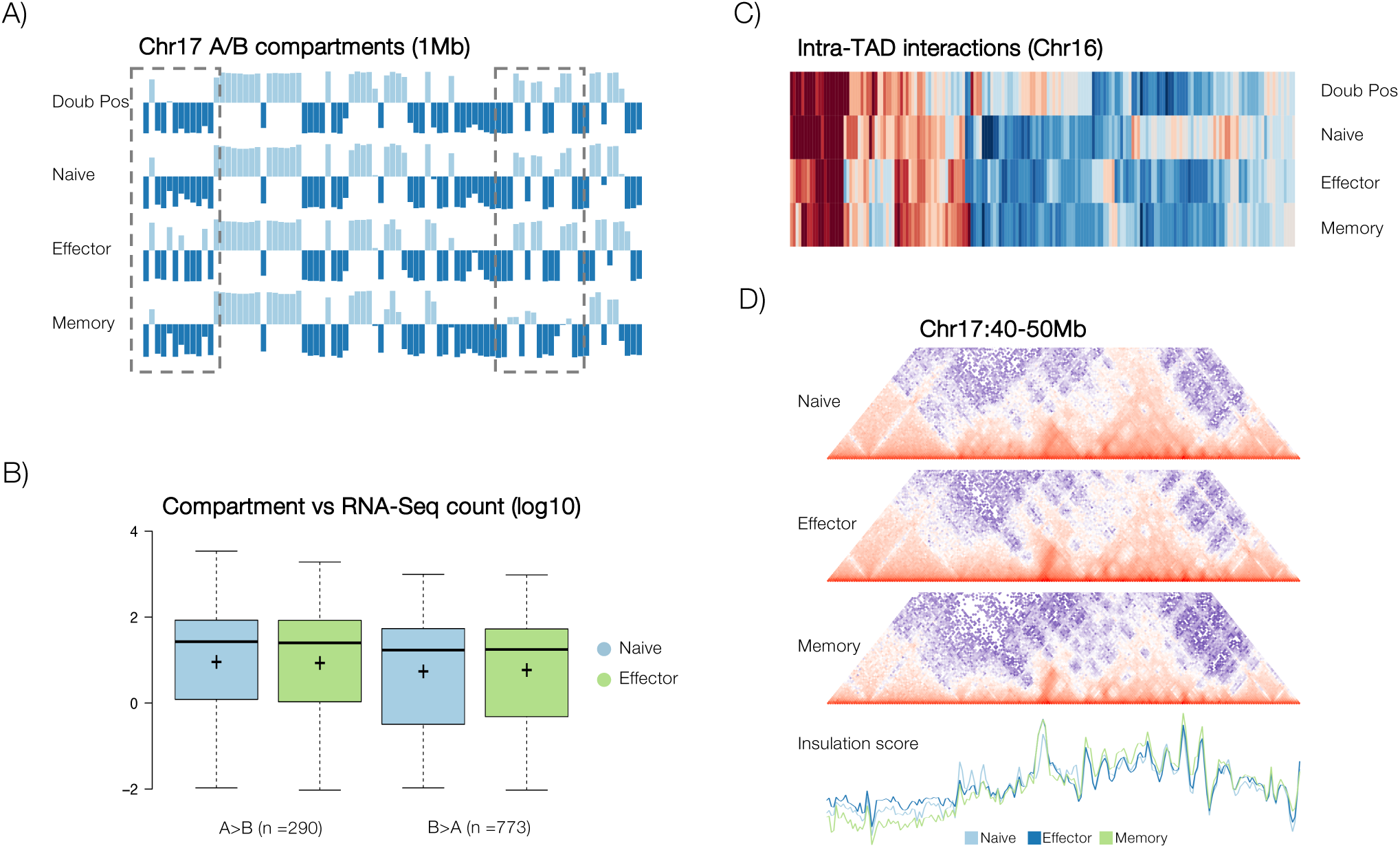
Conservation of higher order chromatin structures during CTL differentiation. Sort purified, naïve (CD44loCD62Lhi) CD45.1^+^ CD8^+^ OT-I CTLs were adoptively transferred into CD45.2^+^ congenic C57BL/6J mice prior to recipients being infected with A/HKx31-OVA. Effector (CD44^hi^ CD62L^lo^) and memory (CD44^hi^) OT-Is were isolated and sort purified either 10 or 60 days p.i., respectively. Virus-specific CTL were compared with sort purified CD4^+^CD8^+^ (double positive) thymocytes from C57BL/6J mice. **A)** Eigenvectors calculated at 1Mb resolution for chromosome 17 of naïve CTL, with A and B compartments shown in light blue and dark blue, respectively. While minor variation in A/B compartment structure was observed between differentiation states (dashed boxes), compartment structures and proportion of the genome in each compartment was largely conserved with differentiation. **B)** Changes in compartment from A to B and B to A with differentiation did not, on average, coincide with changes in gene transcription. Changes in A/B compartment upon naïve (blue) to effector (green) differentiation versus average transcript frequency (log counts per million – cpm) are shown as an example. **C)** Heatmap showing intra-TAD interaction intensities for chromosome 17. **D)** Heatmaps showing interaction frequency within a 10Mb window of chr17, for naïve, effector and memory CTL, with insulation scores calculated for each at 50Kb resolution.

Topological associated domains (TADs) are large scale genomic structures that are largely invariant across cell types and species (Battulin et al., 2015; Dixon et al., 2015; Dixon et al., 2012; Harmston et al., 2017). While the TAD structures are largely invariable, they may switch between A/B compartments to regulate transcriptional activity of genes located within the TAD (Dixon et al., 2015). To examine TAD structures and dynamics during CD8^+^ T cell differentiation, we identified TADs at 1Mb resolution (Cresswell et al., 2020) (see methods), finding that the number (2937 – DP, 2873 – naïve, 2715 – effector, 2923 – memory) and mean size of TADs was similar between differentiation states (**Figures S1D and S1E**). However, while TAD numbers did not vary significantly between differentiation states, intra-TAD interaction frequencies were far more variable suggesting that regulation of chromatin interactions within TADs may underscore distinct CTL states (**Figure 1C**). As an independent means of assessing TAD structure and conservation, Insulation Scores (IS) (Crane et al., 2015) were calculated at 50Kb resolution and compared between differentiation states (**Figure 1D**). We found a strong overlap between samples in the position of local IS peaks, which indicate the position of TAD boundaries, supporting the conclusion that TAD structures are largely conserved across CTL differentiation states. However, importantly, we found that ISs for regions occurring between boundaries were far more variable, again suggesting that differences in interactions occurring within TADs rather than the positioning of TADs or A/B compartments was more likely to explain differences in gene transcription between CTL states.

### CD8^+^ T cell differentiation is associated with intra-TAD reorganisation

Our analysis of higher order chromatin structures indicated that alterations to genomic looping within TAD boundaries was likely a defining feature of CD8^+^ T cell differentiation. To examine this in more depth, a multidimensional scaling analysis (MDS) was performed, comparing *cis* interaction frequencies within 50kb bins between samples. As well as showing a close grouping of biological replicates, this analysis grouped samples into three main clusters; DP and naïve, which clustered separately from all other samples, and separately from each other, and effector and memory, which clustered closely to one another, suggesting a similar genome organisation (**Figure 2A**). Thus, these data again indicated that maturation from DP to naïve, and subsequent differentiation after antigen experience coincided with significant intra-TAD rearrangements of the T cell genome, rather that wholesale changes in genome organisation.

**Figure 2.**
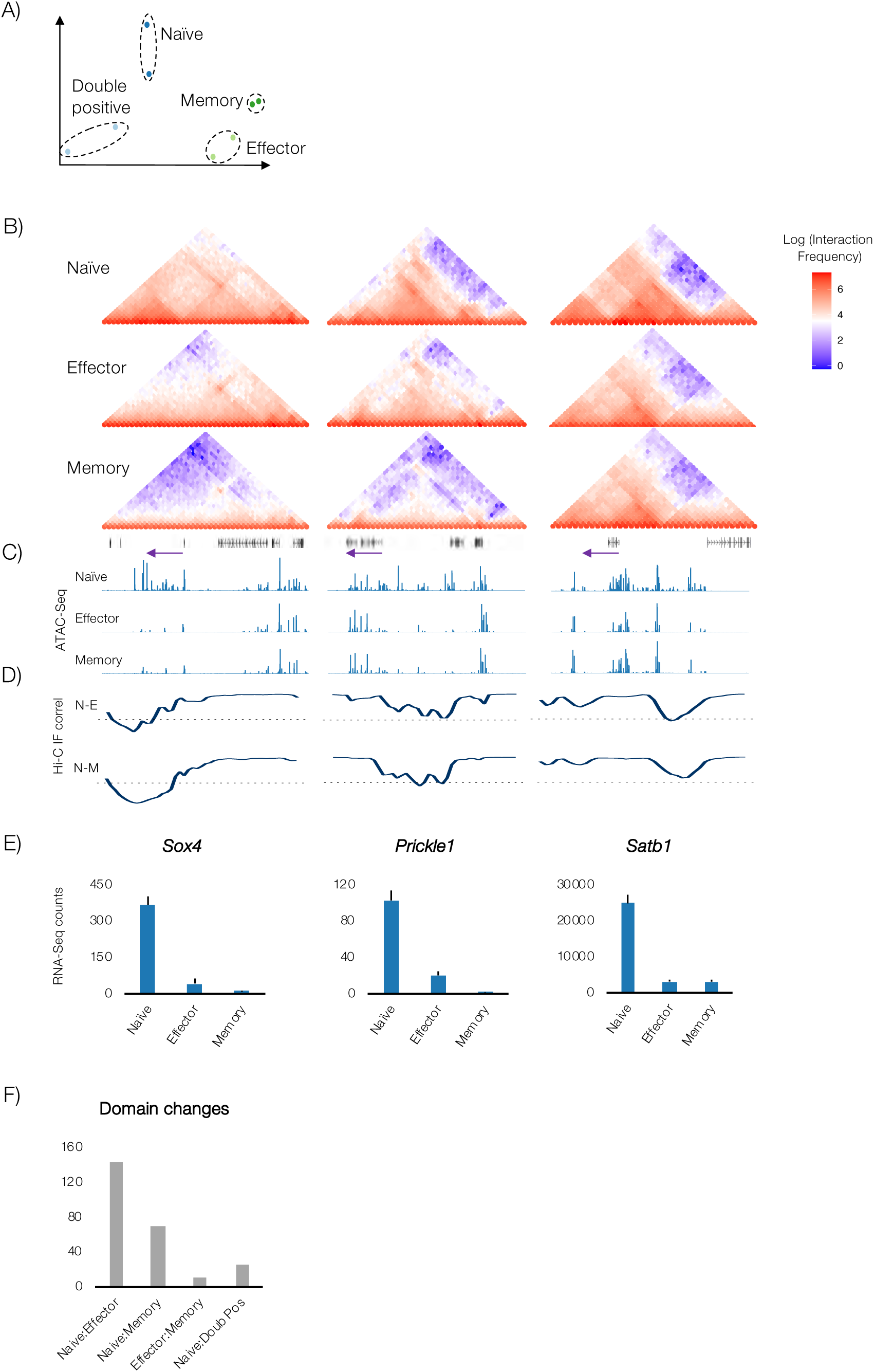
Distinct higher order chromatin structures within distinct CD8+ T cell populations. **A)** MDS plot showing relationship between Hi-C samples derived from double positive (CD4^+^CD8^+^) thymocytes and naïve, effector and memory OT-1 CTLs. **B)** Hi-C data (50Kb bins) normalised using ICED method showing interaction frequency at *Sox4*, *Prickle1* and *Satb1* loci in naïve, effector and memory OT-1 CTLs. Track below memory panel shows genes, with purple arrow highlighting genes of interest and their direction of transcription. **C)** ATAC-Seq data for Naïve, effector and memory OT-1 CTLs. **D)** Pairwise correlation of binned interaction frequencies (50kb) for naïve and effector (N-E), and naïve and memory (N-M) samples, with dotted line indicating 0 on the y axis. **E)** Bottom panel shows normalised RNA- Seq counts (Russ *et al*, 2014) **F)** Quantification of domain changes identified by pairwise correlation (50Kb) analysis depicted in **D.**

As differences identified in the MDS analysis were based on a small bin size (50kb), we reasoned that localised changes in interaction were driving the separation of samples observed, consistent with our finding of differences in interaction frequency and IS occurring largely within boundary elements (**Figures 1B and C**). To identify genomic regions underscoring these differences, we generated heatmaps that identified varying interaction frequencies within naïve, effector and memory CD8^+^ T cell differentiation states and overlaid this with matched ATAC-seq data to measure changes in chromatin accessibility (**Figures 2B and 2C; Figures S3A and S3B**). Differences in Hi-C IFs were identified by calculating pairwise correlations between naïve, effector and memory CD8^+^ T cell states using 50kb bins (**Figure 2D; Figure S2C**). While we found that most bins showed strongly correlated IFs between states, we identified a number of domains that exhibited structural changes visible as large-scale losses (**Figures 2B, 2C, and 2D**) and gains (**Figures 3A, 3B, and 3C**) of IF across broad regions of the genome. Importantly, these changes in contact frequency were also associated with changes in chromatin accessibility and gene transcription (**Figures 2C and 2E; Figures S3B, S3C and S3D**). For instance, loss of IF at loci encoding *Sox4*, *Prickle1* and *Satb1* occurred upon differentiation of naïve CD8^+^ T cells to effector and memory states, and was associated with loss of chromatin accessibility and gene transcription, while gain of IF at loci such as *Prdm1* (encoding BLIMP1) and *Dmrta1* was associated with increased chromatin accessibility and gene transcription (**Figure S3**). Other genes occurring within regions that gained interaction frequency included genes involved in tolerance/co-stimulation including *Cd86*, *Icos*, and *Cblb*, and the killer like receptors *Klra1*, *Klra2*, and *Klrg1*, while examples of genes within regions that lost interaction frequency include *Sox5*, and *Tgfbr2* (**Table S2**). In total, we found the greatest number of domain changes between naïve and effector (144) and naïve and memory (69) samples, with relatively few gross differences separating effector and memory (11) (**Figure 2F**). Interestingly, and suggesting the scale of the differences between the naïve genome architecture and that of effector and memory, naïve and DP were separated by considerably fewer (25) gross changes than naïve and effector or naïve and memory. Again, these data are consistent with the close clustering of effector and memory in our MDS analysis (**Figure 2A**) suggesting a gross change in genome structure following antigen exposure, which is maintained into CTL memory.

**Figure 3.**
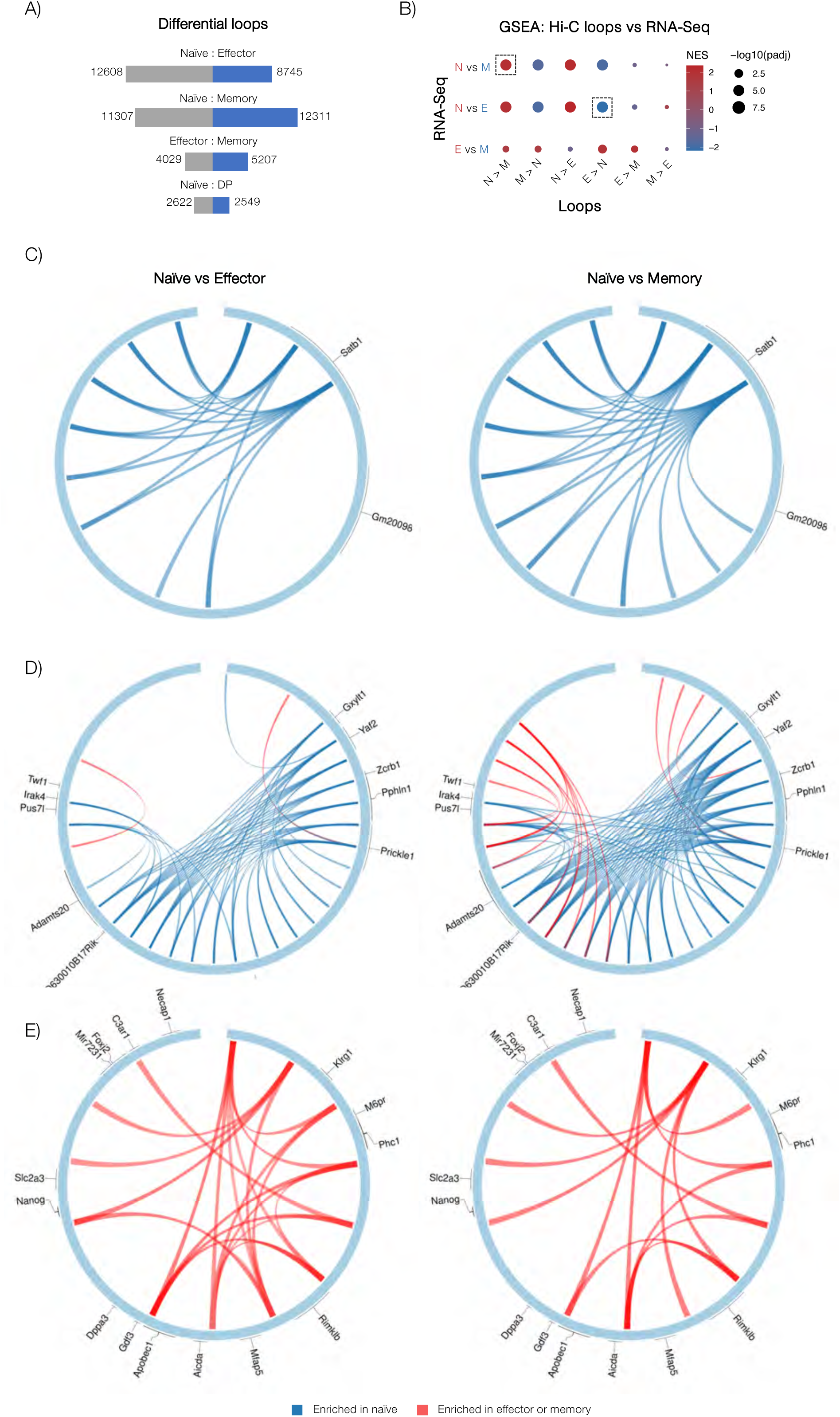
Loss and gain of *cis* regulatory interactions underscores CTL differentiation state specific gene transcription profiles. **A)** Numbers of *cis* interactions unique to each differentiation state, determined by pairwise comparisons using multiHiCcompare (50Kb resolution, 0.05 FDR). **B)** GSEA analysis comparing genes connected by loops enriched in one condition over another (Y axis), against RNA-seq data derived from matching samples (Russ et al., 2014). Circle sizes reflect adjusted p values (-log10) and colour represents normalised enrichment score (NES), with red indicating enrichment versus the first RNA-seq condition listed in pairwise comparison, and blue indicating enrichment is the second RNA-seq condition listed. **C-E**) Examples of loci where loops were lost and gained upon differentiation (blue loops are present in naïve over effector or memory; red loops are gained on differentiation).

### Chromatin looping dynamics underscore CTL differentiation states

To further understand how looping dynamics influence CTL gene transcription following virus infection, we identified loops that were lost or gained following infection using the MultiHiCcompare package (Stansfield et al., 2019). We identified between 5,171 and 23,618 differential loops when samples were compared pairwise, with the largest number separating naïve from memory (23,618), and naïve from effector (21,353), while effector and memory were separated by considerably fewer differences (9,416), again suggesting that these states share a similar genome organisation (**Figure 3A;** see **Table S3** for loops called and gene assignments). Interestingly, naïve and DP samples had fewer differences (5,171) than naïve and effector, or naïve and memory, consistent with our MDS data (**Figure 2A**). Next we assigned loops to the nearest gene (methods) and performed Gene Set Enrichment Analysis (GSEA) to determine whether loss and gain of loops was associated with changes in gene transcription (**Figure 3B**). Overall, we found a strong correspondence between differentiation state specific loss and gain of Hi-C contacts and a corresponding loss and gain of gene transcription. For example, we found that loops enriched in naïve over effector CTLs, and naïve over memory CTLs were associated with genes transcribed more strongly by naïve than effector CTLs (Normalised Enrichment Score (NES) = 2.36), and naïve than memory CTLs, respectively (NES = 2.08; relevant comparisons indicated by dashed boxes). This pattern of altered chromatin interactions tracking with changes in gene expression was also confirmed in the accompanying paper (Quon et al., 2022) using a different infection model and effector subsets, suggesting that this phenomenon does not depend on infection type and reflects intrinsic mechanisms associated with CD8^+^ T cell differentiation programs.

Next we inspected individual gene loci to further understand how fine-scale looping dynamics reflected gene transcription, finding that broadly, loss and gain of looping corresponded with loss and gain of gene expression, respectively. For instance, loci encoding *Satb1*, *Prickle1* and *Sox4* - genes associated with maintenance of stemness and quiescence, which are strongly downregulated following naïve CTL differentiation (**Figure 2E**) – have dense looping structures in naïve CTLs, that are lost on differentiation to effector or memory (**Figures 3B and 3C**; **Figure S3A;** blue ribbons indicate loops present in naïve over effector (left panels), or naïve over memory (right panels), while red ribbons are gained following differentiation of naïve CTLs into effector or memory states). Conversely, loci encoding genes that are expressed following CTL differentiation (*Klrg1* – top panel; *GzmA* and *GzmK* – middle panel; and *Ccl3*, *Ccl4*, *Ccl5*, *Ccl6*, and *Ccl9* – bottom panel; **Figure 3D; Figure S3B**) are characterised by increased looping following differentiation of naïve CTL to effector or memory. Noticeably, looping dynamics following differentiation of naïve CTLs to effector where largely shared with those following differentiation of naïve CTLs to memory, suggesting a mechanism for the rapid recall of effector function following reactivation of memory CTLs, and this was consistent with the close grouping of effector and memory states in our MDS plot (**Figure 2A**).

### Chromatin loops are enriched for differentiation state specific transcriptional enhancers

We next aimed to understand the mechanisms by which differentiation state specific chromatin loops impart transcriptional programs characteristic of the different CD8^+^ T cell states. To do this, we examined the chromatin environment within loops found in naïve but not effector T cells, and vice versa. We found that naïve chromatin loops were enriched for accessible chromatin (measured by ATAC-seq), but the same regions in effector CTLs were not (**Figure 4A**), while effector chromatin loops were enriched for open chromatin in effector CTLs, but the same regions in naïve T cells were not. Moreover, accessible chromatin enrichment patterns for the same regions in memory T cells were very similar to those found for effector T cells, again suggesting that a similar chromatin structure underscores the capacity of memory CTL to elicit effector functions rapidly. Providing a potential explanation for the active chromatin landscape within differentiation state specific chromatin loops, we found that naïve loops were enriched for active and poised enhancers (H3K4me1^+^ H3K4me2^+^; (Russ et al., 2017)) that are present in naïve but not effector T cells, and vice versa (**Figure 4B)**. Indeed, inspecting individual loci, we found that the looping interactions we observed largely connected regions of the genome that were decorated with chromatin features characteristic of active and poised regulatory elements **(Figures 4C and 4D**), including H3K4me1, H3K4me2, H3K4me3, and chromatin accessibility as measured by ATAC-seq (dark blue, light green, dark green, and pink tracks, respectively), although interestingly, some loops did not appear to connect obvious regulatory regions. Moreover, while interactions between putative enhancers and gene promoters were common, we also observed promoter-promoter interactions as reported previously for human CD4^+^ T cells (for instance, at the *Sox4* locus; **Figure S3A**) (Chepelev et al., 2012). Finally, TF enrichment based on curated publicly available lymphocyte ChIP-Seq datasets (Zheng et al., 2019) showed that naïve T cell specific enhancers found within naïve-specific chromatin loops are enriched for binding of TFs including TCF1 and FOXO1, which have roles in maintenance of T cell stemness and quiescence (Danilo et al., 2018; Delpoux et al., 2021; Kerdiles et al., 2009). By contrast, effector specific enhancers found within effector-specific chromatin loops were enriched for binding of TFs such as TBX21 (TBET), IRF4, and PRDM1 (BLIMP1) which have roles in terminal effector differentiation (Intlekofer et al., 2005; Kallies et al., 2009; Man et al., 2013; Rutishauser et al., 2009). Taken together, these data suggest that the dynamics of *cis* regulatory interactions underscores instillation of differentiation specific transcriptional programs within CTLs, largely by connecting genes with enhancers bound by key TFs.

**Figure 4.**
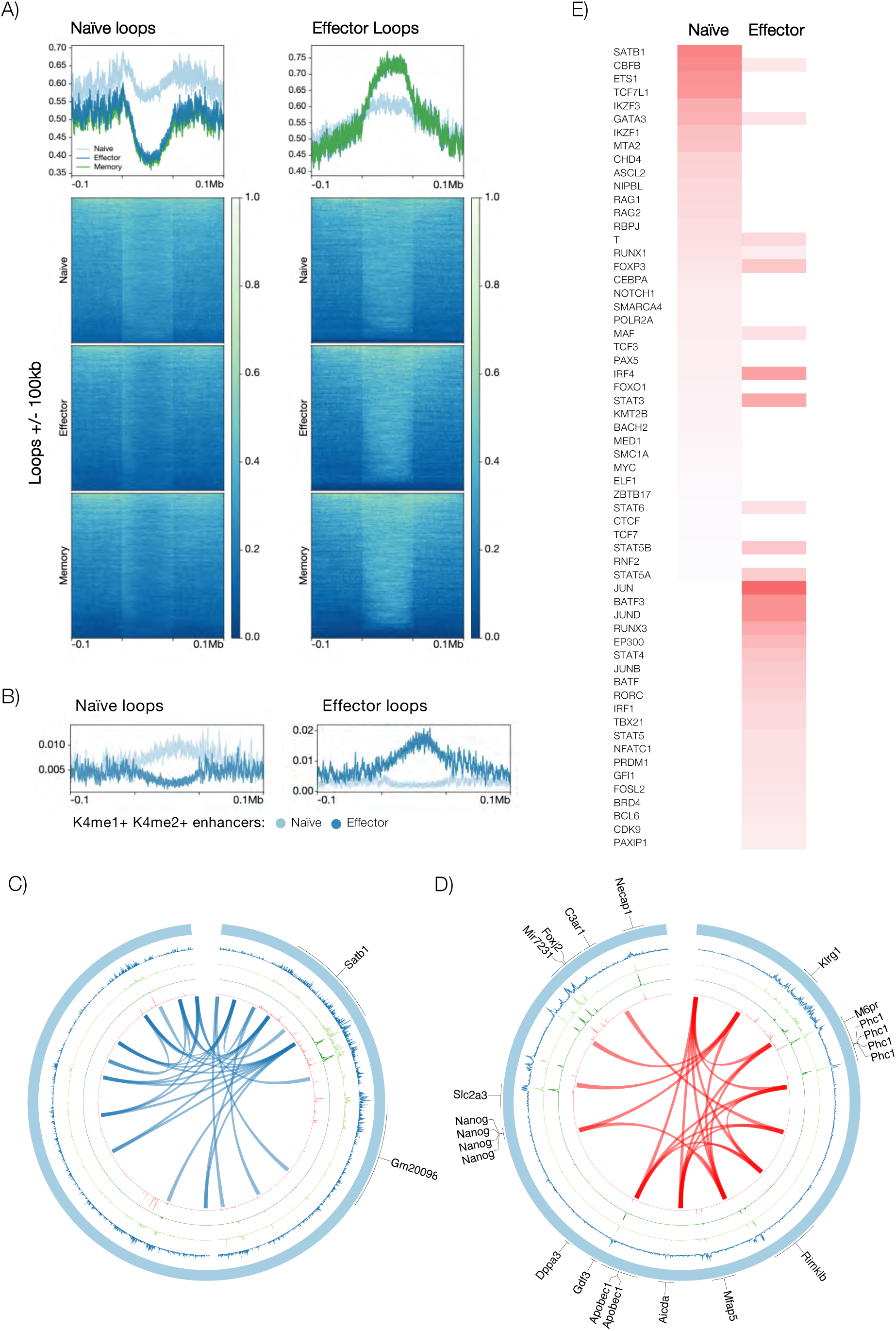
Hi-C Loops border active regions containing differentiation state specific enhancers. **A)** ATAC-seq signal (log2) within and surrounding loops that are present in naïve but not effector CTL, or vice versa. Loops are scaled to occupy 100kb, and ATAC-seq signal is shown for 100kb up and downstream of the loop borders. **B)** Enrichment of active and poised (H3K4me1^+^ H3K4me2^+^) transcriptional enhancers that occur in naive CD8^+^ T cells but not effector T cells within loops that occur in naive but not effector CD8^+^ T cells (upper panel) and vice versa (lower panel). **C, D)** *Cis* interactions connect gene regulatory elements. Circos plots show the gene neighbourhood of *Satb1* **(C)** and *Klrg1* **(D)** in naïve and effector OT-1 CTLs, respectively. Tracks in order from outside to the centre: are genes, H3K4me1 (blue), H3K4me2 (lite green), H3K4me3 (dark green), ATAC-Seq (pink), Hi-C interactions (naïve over effector CTLs **(B)** and effector over naïve CTLs **(C)**) shown as ribbons. **E)** Enrichment of transcription factor binding at TEs unique to naïve or effector (Russ et al., 2017) was performed using curated transcription factor ChIP-Seq data through the CistromeDB Toolbox (Zheng et al., 2019).

### BAHC2 enforces the naïve T cell-specific looping architecture

BACH2 and SATB1 TFBS were enriched within naïve specific chromatin loops (**Figure 4E**). To understand if and how these TFs impact the looping architecture of naïve CTLs, *in situ* HI-C has performed on sort purified naïve CD8^+^ T cells from mice with a point mutation in the DNA binding domain of SATB1 that abrogates DNA binding (Koay et al., 2019) (*Satb1^m1Anu/m1Anu^*) or with BACH2 deficiency (*Bach2^fl/fl^*x *Cd4 Cre* mice; *Bach2^-^*^/-^) (library statistics in **Table S1)**.

To broadly assess changes in genome architecture, an MDS analysis was performed (as in **Figure 2A**), comparing these datasets with the WT naïve, effector and memory datasets described above (**Figure 5A**). We found that while the *Satb1^m1Anu/m1Anu^* datasets overlapped the naïve WT, the *Bach2^-/-^* datasets clustered more closely with effector, suggesting that deletion of BACH2 is sufficient to license the higher order chromatin rearrangement of the naïve genome architecture that accompany effector CTL differentiation. Consistent with this, a pairwise comparison of loops lost and gained between naïve WT and *Bach2^-/-^* identified ∼17,583 differences (**Figure 5B**), which was similar to the number separating naïve and effector (∼21,000; **Figure 3A**), while far fewer loops (4,249) separated effector and *Bach2^-/-^*. Indeed, GSEA analysis showed that genes associated with loops gained in naïve *Bach2^-/-^* CTLs relative to WT naïve, are more highly transcribed in effector than (WT) naïve CTLs (NES 2.79) (**Figure 5C**). Noticeably, the accompanying paper (Quon et al., 2022) found that chromatin changes induced by *Bach2* deletion was more similar to the activated T cell subsets with greater differentiation potential (MP and memory). Thus, deletion of *Bach2* shifts the chromatin architecture to resemble the more stem-like antigen-experienced CD8^+^ T cell subsets.

**Figure 5.**
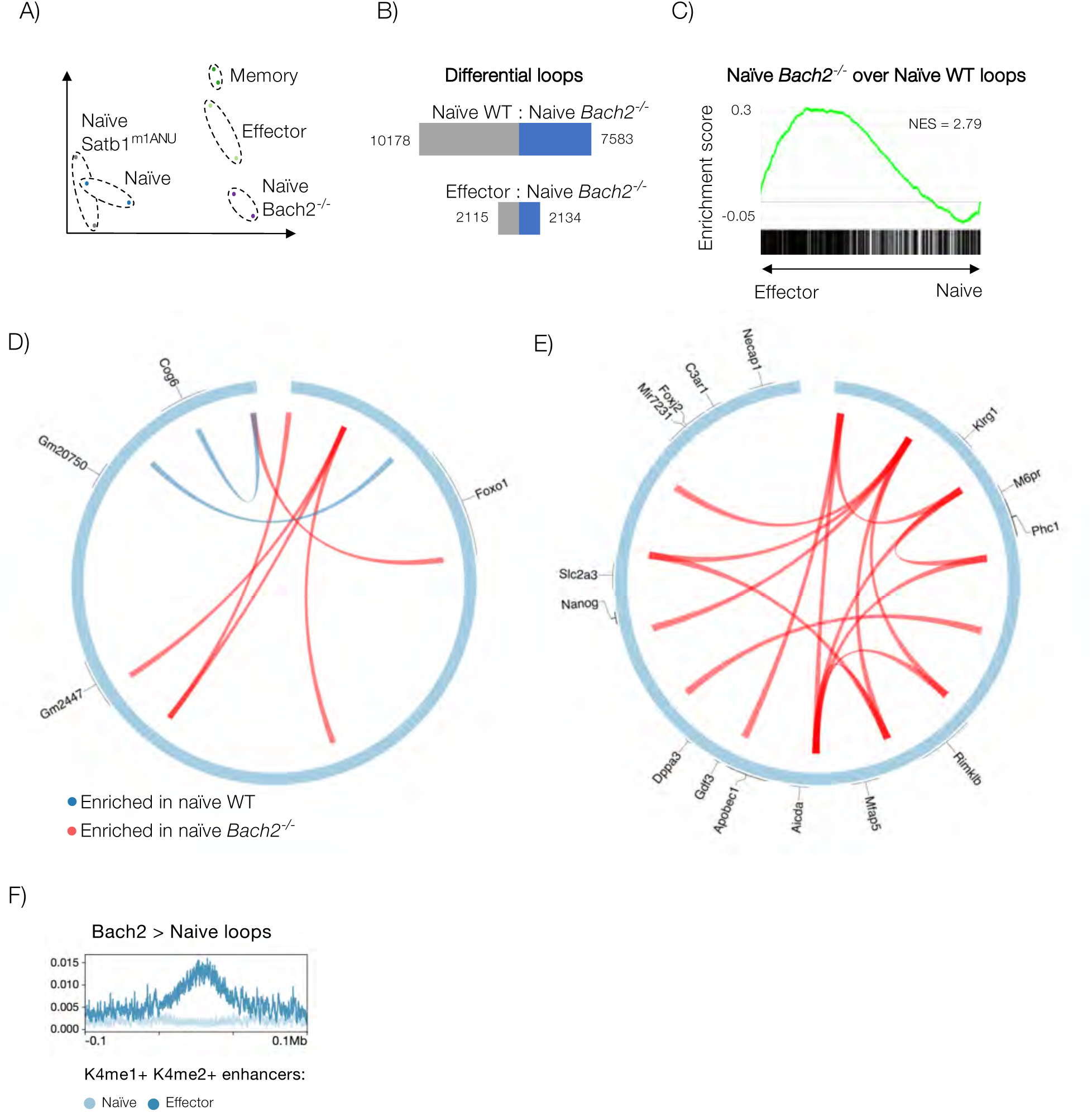
BACH2 enforces a naïve chromatin architecture. (**A)** MDS plot showing relationship between naïve *Bach2*^-/-^ and naïve *Satb1^m1Anu/m1Anu^*(described in figure 7) Hi-C samples and naïve, effector and memory OT-1 CTLs. (**B)** Loss and gain of *cis* interaction in naïve *Bach2^-/-^* CD8^+^ T cells in comparison with WT naïve and virus specific OT-1 CD8^+^ T cells. **C)** GSEA analysis comparing genes connected by loops gained in naïve *Bach2^-/-^* CD8^+^ T cells relative to naïve WT CD8^+^ T cells against RNA-seq data derived from naïve and effector CTLs samples (Russ et al., 2014). P values and normalised enrichment score (NES) are shown. **(D**, **E)** Examples of changes in looping architecture in naïve *Bach2^-/-^* CD8^+^ T cells relative to naïve WT CD8^+^ T cells (blue loops are enriched in WT naïve over naïve *Bach2^-/-^* CD8^+^ T cells and red loops are enriched in naïve *Bach2^-/^*^-^ CD8^+^ T cells over WT). **(F)** Loops that occur in naïve *Bach2^-/-^* CD8^+^ T cells but not WT naive CD8^+^ T cells are enriched for active and poised (H3K4me1^+^ H3K4me2^+^) transcriptional enhancers that occur in effector CD8^+^ T cells but not naïve T cells.

Finally, we inspected individual gene loci to further understand how deletion of *Bach2* altered looping dynamics. Consistent with BACH2 having a role in enforcing naïvety, we found that the locus encoding *Foxo1*, which is itself required to enforce CTL naïvety (Delpoux et al., 2021) was reorganised in *Bach2^-/-^* CTLs, with both loss (blue) and gain (red) of loops relative to the WT (**Figure 5D)**. Indeed, *Bach2^-/-^*CTLs showed loss of a loop connecting the *Foxo1* promoter and a downstream non-coding element, suggesting that BACH2 maintains CTL naïvety in part by driving FOXO1 expression. Importantly, we also found a loss of looping at the *Tcf7* and *Lef1* loci, and gain of loops at the *Prdm1* (BLIMP1), *Tbx21* (TBET)*, Zeb2*, *Nfatc4* encoding loci, suggesting a loss of naïve potential and engagement of the effector CTL transcriptional program (**Table S3**). Consistent with this, the *Klrg1* encoding locus also underwent large-scale reorganisation in *Bach2^-/-^*CTLs, although the dynamic was different to that at the *Foxo1* locus, with loops being gained but not lost in *Bach2^-/-^* CTLs (**Figure 5E)**. Suggesting that BACH2 restrains a largely autonomous effector differentiation program that is engaged following CTL activation, loops gained at the *Klrg1* locus with deletion of *Bach2* where largely identical to those gained on differentiation of WT naïve to effector and memory following virus infection (**Figure 3E**). Moreover, loops that were acquired in *Bach2^-/-^* T cells occurred in regions that harbour enhancers that are active and poised (H3K4me1^+^ H3K4me2^+^) in effector but not naïve CD8^+^ T cells. Further, the accompanying paper (Quon et al., 2022) found an enrichment of CTCF binding upstream of regions with altered chromatin interactions with *Bach2* deletion, suggesting that BACH2 may collaborate with CTCF to regulate the chromatin interactions. Thus, taken together, these data indicate that BACH2 is essential for maintenance of CTL naïvety because it enforces a looping architecture that maintains naïve T cell quiescence and stemness functions, while blocking engagement of effector transcriptional programs.

### A distal role for SATB1 in maintenance of a naïve-specific looping architecture

Given SATB1 binding sites were enriched within naïve-specific enhancers (**Figure 4E**), and our recent data demonstrating that naïve *Satb1^m1Anu/m1Anu^* CD8^+^ T cells have an activated phenotype (Nussing *et al*. *in press*), this suggested that SATB1 may also play a role in ensuring CTL naïvety via chromatin organisation. To understand this further, a pairwise comparison of looping architectures was performed on WT naïve and *Satb1^m1Anu/m1Anu^* naïve Hi-C datasets described in **Figure 5A**. We found 562 differential loops in this comparison (**Figure 6A; Table S3**), much fewer than that described for *Bach2^-/-^* (**Figure 5B**) and comparisons between virus-specific OT-1 CTL datasets (**Figure 3A**). Next, we performed GSEA analysis to determine whether changes in loop architecture might underscore the altered phenotype and transcriptome of *Satb1^m1Anu/m1Anu^*mice. To do this we compared genes associated with loops gained or strengthened in naïve *Satb1^m1Anu/m1Anu^* CTL (relative to the WT), with RNA-seq data from naïve C57BL/6J and naïve *Satb1 ^m1Anu/m1Anu^* CTLs (**Figure 6B**) (Nussing *et al*. *in press*). Indeed, we found that genes associated with *Satb1^m1Anu/m1Anu^* specific loops tended to be upregulated in *Satb1 ^m1Anu/m1Anu^* CTLs over the WT, indicating that the altered looping architecture was likely driving the activated transcriptome and phenotype of *Satb1^m1Anu/m1Anu^* CTLs.

**Figure 6.**
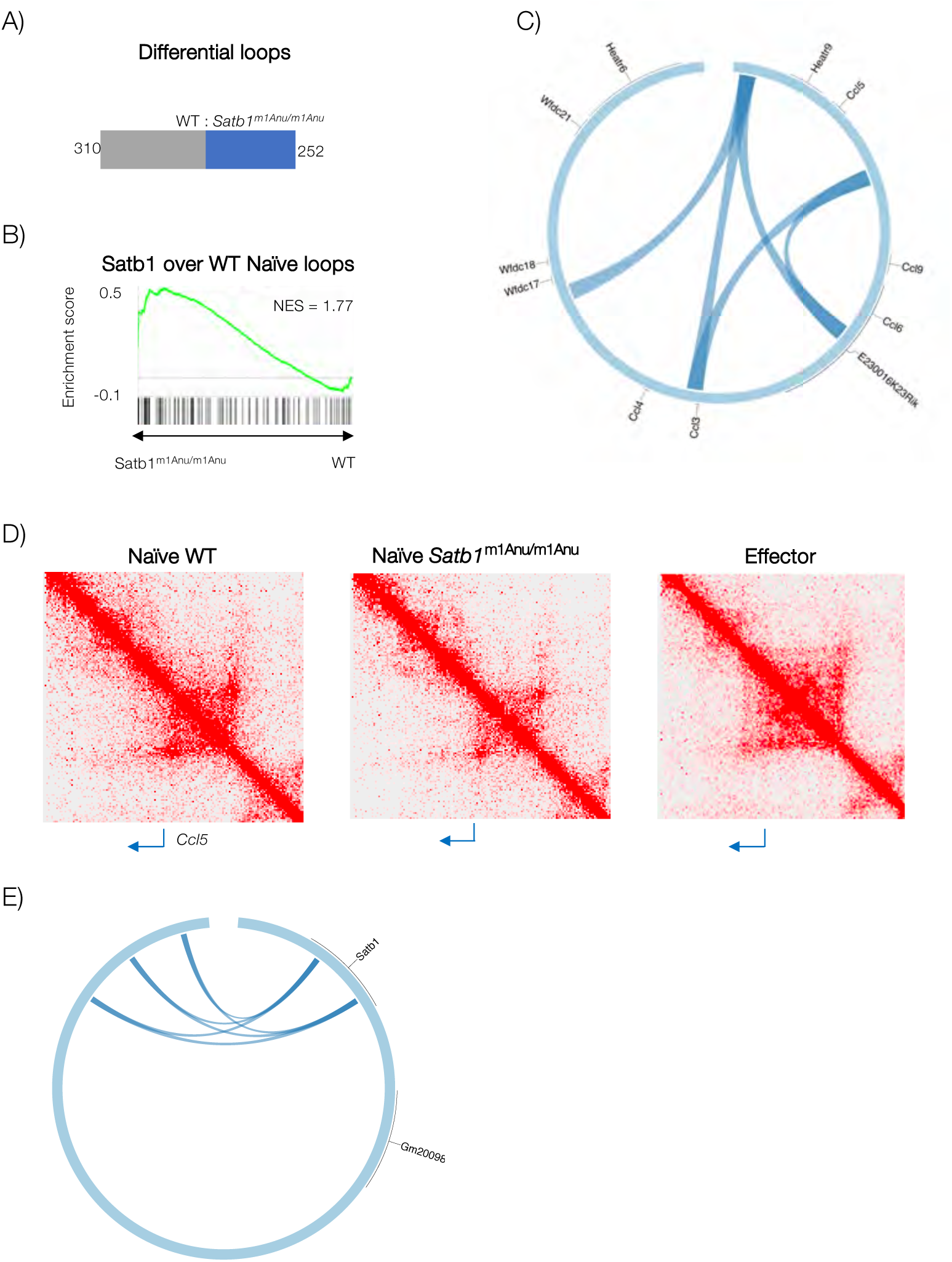
SATB1 maintains CD8^+^ T cell naïve chromatin architecture. **(A)** Loss and gain of cis interactions in naïve *Satb1^m1Anu/m1Anu^* CD8^+^ T cells in comparison with WT naïve OT-1 T cells. **(B)** GSEA analysis comparing genes connected by loops gained in naïve *Satb1*^m1Anu/m1Anu^ CD8^+^ T cells relative to naïve OT-1 T cells against RNA-seq data derived from matching samples (Nussing et al. *under revision*). Normalised enrichment score (NES) is shown. **(C)** Loops lost at the type 1 chemokine locus in naïve *Satb1*^m1Anu/m1Anu^ CD8^+^ T cells relative to naïve OT-1 cells. **(D)** Hi-C contact maps showing the *Ccl5* encoding locus in naïve OT-1, *Satb1^m1Anu/m1Anu^*naïve, and effector OT-1 CTLs. **(E)** Loops lost at the *Satb1* locus in naïve *Bach2^-/-^* CD8^+^ T cells relative to naïve OT-1 cells.

Next, we performed GSEA analysis to determine whether changes in loop architecture might underscore the altered phenotype and transcriptome of *Satb1^m1Anu/m1Anu^* mice. To do this we compared genes associated with loops gained or strengthened in naïve *Satb1^m1Anu/m1Anu^* CTL (relative to the WT), with RNA-seq data from naïve C57BL/6J and naïve *Satb1 ^m1Anu/m1Anu^* CTLs (**Figure 6B**) (Nussing *et al*. *in press*). Indeed, we found that genes associated with *Satb1^m1Anu/m1Anu^* specific loops tended to be upregulated in *Satb1 ^m1Anu/m1Anu^* CTLs over the WT, indicating that the altered looping architecture was likely driving the activated transcriptome and phenotype of *Satb1^m1Anu/m1Anu^* CTLs. To understand this further, we inspected the dynamics of looping loss and gain at individual gene loci. We found that at the type 1 chemokine locus (encoding *Ccl3*, *Ccl4*, *Ccl5*, *Ccl6* and *Ccl9*), there was a loss of loops in the *Satb1^m1Anu/m1Anu^* CTLs, relative to the WT, despite *Ccl5* being upregulated in the former (**Figure 6C**). Moreover, closer inspection of contact matrices confirmed a partial loss of contact frequency across the region in the *Satb1^m1Anu/m1Anu^* CTLs, which also appeared to occur in effector CTLs together with a “spreading” of the zone of contacts (**Figure 6D**). Thus, it appeared that acquisition of *Ccl5* transcription within effector CTLs required a gross and stepwise remodelling of looping architecture, with *Satb1^m1Anu/m1Anu^*CTLs having an architecture and transcriptional profile intermediate between WT naïve and effector.

Finally, fewer alterations to the *Satb1^m1Anu/m1Anu^* looping architecture relative to that observed in the *Bach2^-/-^* dataset (**Figures 6A** and **5b**, respectively) suggested that SATB1 plays a role distal to BACH2. Indeed, inspection of the looping architecture at the *Satb1* locus in the naïve *Bach2^-/-^* dataset showed a partial loss of loops present in naïve CD8^+^ T cells that are lost upon differentiation to effector (**Figures 6E** and **3C**, respectively), while the architecture of the *Bach2* locus was not altered in the *Satb1^m1Anu/m1Anu^* dataset. Thus, taken together, these data indicate that SATB1 maintains CTL quiescence through enforcement of a naïve chromatin looping architecture that appears to be downstream of BACH2.

### Altered chemokine expression in mice following deletion of *cis* interacting elements mapped by Hi-C

Having found that the dynamics of loss and gain of *cis* interactions broadly described the installation and maintenance of CTL differentiation state specific transcriptional programs (**Figures 3A and 3B**), we next asked whether these interactions were necessary drivers of those programs. We had previously utilised ChIP-seq to identify putative transcriptional enhancers of *Ccl5* (Russ et al., 2017) located at −5Kb and −20Kb region upstream of the *Ccl5* promoter (Russ et al., 2017). Given SATB1 mutation altered chromatin looping at the *Ccl5* locus (**Fig. 6**), we performed virtual chromosome confirmation capture (4C) at 5Kb resolution, anchored at the *Ccl5* TSS to determine if there is altered chromatin looping was evident between naïve and effector CD8^+^ T cells. Our Hi-C data demonstrated an increased interaction frequency between the *Ccl5* promoter and a region spanning ∼20Kb upstream (**Figure 7A**) which correlated with the −5 and −20Kb *Ccl5* enhancers. To validate the virtual 4C analysis, we carried out Formaldehyde Assisted Isolation of Regulatory Elements (FAIRE) to assess changes in chromatin accessibility associated with CD8^+^ activation (**Fig. 7B**). This indicated that the increase in interaction frequency observed within the Hi-C data, correlated with an increase in chromatin accessibility at the −5Kb and −20Kb *Ccl5* enhancers, indicative of transcriptional licensing of these promoters (Russ et al., 2017). To assess the functional impact of these regulatory enhancers on *Ccl5* transcription, we utilised CRISPR/Cas9 genome targeting to generate two separate mouse lines with deletions at the *Ccl5* −5Kb or −20Kb enhancers (Δ−5Kb and Δ−20Kb lines, herein; **Figure 7B**). The Δ−5Kb and Δ−20Kb lines and wild-type (WT) C57BL/6 controls were infected intranasally with Influenza A/HKx31 virus, and lymphocytes from bronchiolar lavage fluid (BAL), spleens and draining lymph node (mediastinal lymph node; MLN) were sampled 10 days post infection, and chemokine expression was assessed by ICS. Further, the body weight of mice was monitored throughout the course of the infection, where we found that both mutant lines lost significantly more weight than the WT, with the Δ−20Kb line having the most significant weight loss (days 3 to 9 post infection; p= <0.01; **Figure S4A).** We found that within each tissue, the Δ−5Kb deletion nearly completely abolished CCL5 production, both by CTLs and CD4^+^ T cells, while surprisingly, the Δ−20Kb deletion did not impact CCL5 expression in either subset, despite this line having the most significant weight loss following influenza challenge **(Figure 7C**; **Figure S4B**). Thus, these data demonstrate that acquisition of the loop connecting the −5 enhancer with the CCL5 promoter is required to enable CCL5 expression within effector CTLs and CD4^+^ T cells.

**Figure 7.**
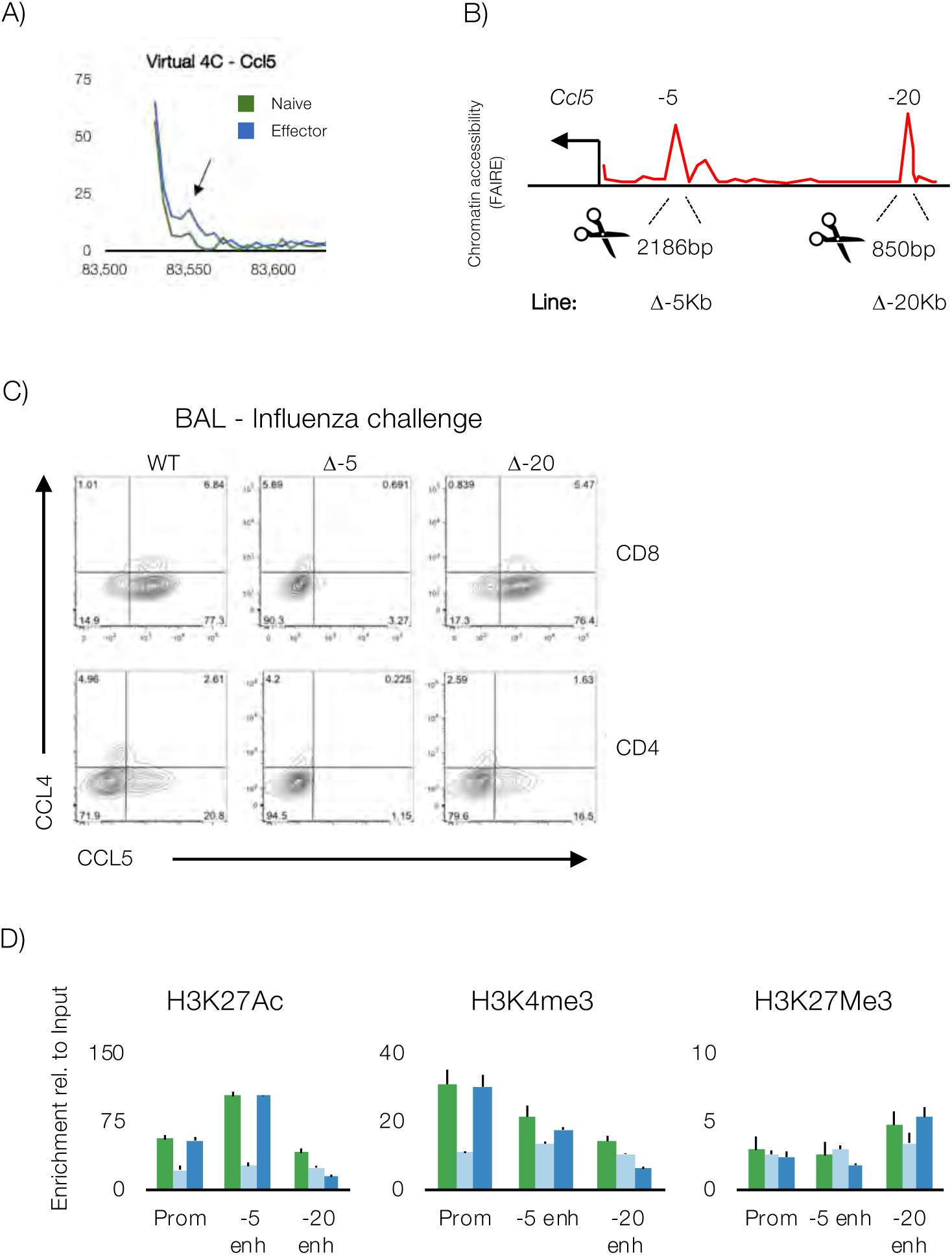
Altered chemokine expression in mice following deletion of cis interacting elements mapped by Hi-C. **(A)** Identification of an interactions between the *Ccl5* gene promoter and previously identified transcriptional enhancers at −5kb and −20kb from the *Ccl5* transcription start site (Russ et al., 2017). Data is presented as a virtual 4C plot, showing naïve and effector Hi-C data, with the arrow indicating a zone of increased interaction in effector CTLs. **(B)** Chromatin accessibility data (mapped by FAIRE) in effector CTL, showing the positioning of CRISPR deletions made in separate mouse lines to remove the −5 and −20 transcriptional enhancers. **(C)** Wild-type and enhancer deletion mice were infected intranasally with 10^4^ pfu A/HKx31influenza virus, and lymphocytes were collected from the bronchiolar lavage (BAL) fluid on d10 for analysis by flow cytometry to assay CCL4 and CCL5 expression in CD8^+^ and CD4^+^ T cells. **(D)** Reduced H3K27Ac at the *Ccl5* locus in in vitro cultured enhancer deletion effector CTLs. Naïve CTL from WT (blue) and −5 (red) and −20 (green) enhancer deletion mice were stimulated with plate bound αCD3 and αCD28 and cultured for 5 days before ChIP assays were performed to measure histone acetylation at the promoter and enhancers of *Ccl5*. Data are pooled from 3 independent cultures, and error bars are SEM. Data are expressed relative to a total input.

To understand the mechanism by which deletion of loop ends impacts CCL5 expression, ChIP was performed on *in vitro* effector CTLs to probe the chromatin composition of regions immediately adjacent to the −5Kb and −20Kb deletions, as well as the *Ccl5* promoter (H3K4me3 and H3K27Ac, which mark active chromatin, and H3K27me3, which marks repressed chromatin; **Figure 7D**). In WT CTLs, we found enrichment of H3K4me3 and H3K27Ac at all 3 regions, while H3K27me3 was distributed evenly across the locus, albeit at low enrichment levels. In contrast, Δ−5Kb CTLs had diminished levels of the permissive modifications across the locus, while Δ−20Kb CTLs had a minor reduction in levels of the permissive modifications specifically adjacent to the deletion site. Thus, these data suggested that deleting loop ends impacts the ability of the *Ccl5* locus to acquire a transcriptionally permissive chromatin following CTL activation, consistent with our finding that differentiation specific loops demarcate regions of open chromatin (**Figure 4A**).

## DISCUSSION

Changes in differentiation state underscore the capacity of CD8^+^ T cells to mediate pathogen clearance and form protective immunological memory, with these state transitions driven by gross transcriptional reprogramming (Kaech et al., 2002; Russ et al., 2014). While regulated enhancer usage (Russ et al., 2014; Scott-Browne et al., 2016; Sen et al., 2016; Yu et al., 2017), TF binding (Intlekofer et al., 2005; Kallies et al., 2009; Man et al., 2013; Rutishauser et al., 2009), and chromatin composition (Barski et al., 2007; Russ et al., 2014) are important determinants of transcriptional reprogramming, how these factors interact within the 3-dimensional space of the nucleus to modulate CD8^+^ T cell differentiation is not well understood. Our data demonstrate that transcriptional reprogramming is choreographed by changes in chromatin looping, with differentiation state-specific loops demarcating regions of open chromatin, within which cell state appropriate enhancers, TFs and genes are brought together to install and maintain transcriptional programs.

TADs serve to partition the genome into discrete functional units that confine regulatory activity to specific chromatin domains (Lieberman-Aiden et al., 2009). Established TADs are generally considered largely invariant across cell types and with cellular differentiation (Battulin et al., 2015; Dixon et al., 2015; Dixon et al., 2012; Harmston et al., 2017). In support of these earlier studies we observed no difference in these higher order chromatin TAD structures between naïve, effector and memory CTL populations. Importantly, CD8^+^ T cell differentiation was associated with discrete changes in the spatial organisation of chromatin looping within TADs. These observations are also consistent with those showing that B activation and subsequent differentiation is associated with discrete changes at the level of chromatin looping, as opposed to higher order genome structures (Johanson et al., 2018). Interestingly, a recent study that examined changes in chromatin looping after polyclonal activation of naïve human T cells reported apparent alterations in TAD boundaries in both activated CD4^+^ and CD8^+^ T cells (Bediaga et al., 2021). These changes in chromatin topology upon activation involved an increase in looping frequency within small domains reminiscent of subTAD structures that have been described (Rao et al., 2014). Whether these represent subTADs, or small de novo TADs is an open question.

A surprising finding of this study was that deletion of *Bach2* resulted in wholesale remodelling of the naïve genome to one which is architecturally similar to that of effector CTLs. This finding implies that CD8^+^ T cell differentiation is a largely autonomous process, with BACH2 maintaining naïvety by repressing the formation of loops that underscore the effector transcriptional program. This finding is consistent with studies showing that BACH2 restrains CTL differentiation (Roychoudhuri et al., 2016; Utzschneider et al., 2020) by competing with AP-1 factors for binding of enhancers within naïve T cells that, once activated, drive the terminal effector differentiation program (Roychoudhuri et al., 2016). Interestingly, AP-1 binding requires phosphorylation of AP-1, which is itself linked to TCR activation. As we observed remodelling in the absence of infection, it remains to be determined precisely how and why BACH2 deficiency licences effector T cell differentiation, and whether or not events subsequent to deletion are required to promote genome remodelling. In either case, these observations demonstrate that loss of BACH2 licences gross remodelling of the naïve genome and loss of CD8^+^ T cell naïvety.

Consistent with the BACH2 data indicating that specific TFs restrain CD8^+^ T cell differentiation by enforcing a naïve chromatin architecture, we found that mutation of SATB1 resulted in a partially reconfigured genome architecture within naïve CD8^+^ T cells, consistent with a previous report of SATB1 as a chromatin organiser in CD4^+^ T cells (Cai et al., 2006). This finding is also consistent with transcriptional data (Nussing et al., *under revision*) demonstrating that naïve CD8^+^ T cells from *Satb1^m1Anu/m1Anu^* mice have a transcriptome showing hallmarks of early activation, with upregulation of genes including *Pdcd1* (encoding PD-1), *Cd44* and *Il2rb*. Interestingly, and in contrast to the *Bach2* mutant, remodelling of the *Satb1^m1Anu/m1Anu^* genome appears to arrest early, suggesting that SATB1 and BACH2 operate at different points of a regulatory hierarchy that maintains the naïve genome architecture. Indeed, the finding that the *Satb1* locus is remodelled in the *Bach2* mutant, but not *vice versa*, and the less dramatic alteration of genome architecture in the *Satb1* mutant suggests that that SATB1 plays a role downstream of BACH2 and may act to fine-tune genome structure.

In line with the concept that higher order chromatin structures within the genome are maintained by key chromatin binding factors, a recent study showed that compound deletion of *Lef1* and *Tcf7* within naïve CD8^+^ T cells also resulted in an altered genome architecture and transcriptome, however, changes occurred across multiple levels of genome organisation, including at the level of compartments, TAD structures, and looping (Shan et al., 2021). Moreover, the transcriptional changes observed in that study included upregulation of effector program genes within naïve cells, but also many non-lineage genes including those normally expressed by other lymphocytes including B cells and NK cells, as well as myeloid lineage cells including granulocytes. While it is not clear whether these TFs mediate chromatin spatial organisation directly, or somehow regulate expression of chromatin organising proteins, these studies highlight that specific TFs not only regulate different aspects of genome structure to maintain T cell naïvety, but also lineage fidelity.

We found that naïve CD8^+^ T cells have a genome architecture that is distinct from either effector or memory, with large-scale architectural changes needed to enact lineage-specific function. By comparison, effector and memory genomes are structurally similar, suggesting that the ability of memory T cells to rapidly recall effector function is in part due to a pre-configured looping architecture, and indeed we found that loops present in effector cells were enriched for open chromatin in resting memory cells (Figure 4a). These data are consistent with previous reports that effector loci within resting memory T cells are characterised by open chromatin, despite not being transcribed (Denton et al., 2011; Northrop et al., 2008; Russ et al., 2014; Zediak et al., 2011). A recent study showed that TCF1 is required to instruct the looping architecture within central memory CD8^+^ T cells, with TCF1 ablation resulting in an inability to engage transcription of genes required for secondary expansion and metabolic reprogramming (Shan et al., 2022). These observations imply that genome architecture is not sufficient to instruct cell type specific gene transcription, but rather, in the case of memory CTLs, TFs also serve to preconfigure the spatial organisation of chromatin to transcriptionally poise appropriate genes for rapid activation following secondary challenge.

Finally, our data highlight a crucial and unique distinction between naive and effector/memory CD8^+^ T cell states, namely the spatial and looping interactions observed within higher order chromatin structures. In particular, our data point to key chromatin binding proteins as providing the molecular restraint that is actively enforced in naive CD8^+^ T cell, which is distinct from the ‘rapid fire’ capacity of effector/memory CD8^+^ T cells. Nevertheless, T cell activation is associated with chromatin remodelling, albeit in a discrete and targeted way. CTCF has been previously implicated in playing a role in the establishment of TAD structures during embryonic cell development (Chen et al., 2019). The accompanying paper (Quon et al., 2022) found that knockdown of CTCF, a known regulator of genome organization, prevented terminal CD8^+^ T cell differentiation by disrupting CTCF binding at weak-affinity binding sites to promote the memory transcriptional program at the expense of the terminal differentiation transcriptional program. A specific CTCF binding site at an effector-specific enhancer in the type I chemokine locus was also identified to insulate CCL3 expression, suggesting that CTCF may be important for regulation of specific enhancer-promoter interactions. Further, depletion of YY1, a protein known to regulate looping within CTCF-mediated chromatin loops (Beagan et al., 2017), also prevented the formation of terminal effector cells. Our data, combined with the associated study (Quon et al, 2022) demonstrate that, not only are specific transcription factors needed to reinforce the chromatin architecture needed for either naïve or effector/memory states, but other chromatin remodelling factors such as CTCF and YY1 are necessary to sustain the terminal differentiation of CD8^+^ T cells.

## ACKNOWLEDGEMENTS

We thank the Monash Flowcore facility (Monash University, Clayton) for helpful advice and technical assistance with flow cytometry and cell sorting experiments, and the Monash Micromon Genomics (Monash University, Clayton) and Hudson Genomics (Hudson Institute, Monash University Health and Medical Precinct, Clayton) facilities for advice relating to preparation and sequencing of Hi-C and ATAC-Seq samples. Bioinformatic analyses were performed by the Monash Bioinformatics Platform, and mice were bred and housed at the Monash Animal Research Platform, at Monash University, Clayton. Thanks to the Goldrath, Turner and La Gruta laboratories for technical advice, helpful discussion, and critical reading of the manuscript. This work was supported by grants from the National Health and Medical Research Council of Australia, APP1003131 (S.J.T); an Australian Research Council Discovery Grant, DP170102020 (S.J.T); a joint Monash University-University of California, San Diego Seed Development grant (S.J.T and A.W.G); an NIH P01AI13212 (A.W.G.) and NIH grant AI102853 (C.M.).

## AUTHOR CONTRIBUTIONS

Conceptualisation, B.E.R., S.Q., B.Y., S.J.T.; Methodology, B.E.R., J.L., K.T.; Investigation, B.E.R., B.Y., J.L., J.K.C., S.N., A.E.M., V.A.U., T.J.B., I.A.P., A.L.S.; Software, K.T.; Formal analysis, B.E.R, K.T., M.O., Z.H., P.F.H., A.B., M.S. D.P.; Resources, A.K.; Writing - Original Draft, B.E.R.; Writing - Reviewing Editing, B.E.R., S.J.T., S.Q., A.W.G., P.C., C. M., A. K; Supervision, S.J.T., A.W.G., D.P., C.M. P.C.; Funding Acquisition, S.J.T. A.W.G., C.M.

## RESOURCE AVAILABILITY

### Lead Contact

Further information and requests for resources and reagents should be directed to and will be fulfilled by the lead contact, Prof Stephen Turner.

### Materials Availability

CCL5 mouse lines are available contingent on signing of appropriate Material Transfer Agreements between Institutions. All other materials are freely available.

### Data and Code Availability

Hi-C and ATAC-seq data have been deposited at GEO and are publicly available using the GEO accession number: GSE225885. Other data accession numbers are listed in the key resources table. All original code is publicly available as of the date of publication. RRIDs are listed in the key resources table. Any additional information required to reanalyse the data reported in this paper is available from the lead contact upon request

## EXPERIMENTAL MODEL AND SUBJECT DETAILS

### Mice

Ly5.2^+^ C57BL/6J, *Satb1*^m1Anu/m1Anu^, and Ly5.1^+^ OT-I mice were bred and housed under specific-pathogen-free conditions at the Monash Animal Research Platform, with housing and experimental procedures approved by the Monash University Animal Ethics Committee. *Bach2^fl/fl^* x *Cd4Cre* mice were bred and housed under specific-pathogen-free conditions at the Department of Microbiology and Immunology Animal Facility at the University of Melbourne. All mice used were female, and aged 8-12 weeks old. For infection, mice were anaesthetised and infected i.n. with 10^4^ p.f.u. of recombinant A/HKx31 virus engineered to express the OVA257–264 peptide (x31-OVA) in the neuraminidase stalk. For adoptive transfer studies, CD45.1^+^ OT-I T cells were adoptively transferred into female CD45.2^+^ recipients.

### Primary Cell Cultures

Naive CD8𝛼+ CD44lo/int cells were sort-purified from C57BL/6J or Δ−5Kb and Δ−20Kb mice (8-12 weeks) (> 99% purity). Cultures were initiated by stimulating 3.3 × 105 T cells with plate-bound anti-CD3ε (10ug/ml), anti-CD28 (5ug/ml), and anti-CD11a (10ug/ml) antibodies, and cultured in the presence of IL-2 (10U/ml). Cells were cultured in 3ml RPMI, supplemented with 10% FCS (v/v), 2mM L-glutamine, and penicillin and streptomycin in 6-well plates, before being expanded into T25 flasks (10mls media) after 72hrs, T75 flasks at 96hrs (20ml media). Cultures were harvested at 120hrs.

## METHOD DETAILS

### ATAC-seq

We used an ATAC-seq protocol adapted from (Buenrostro et al., 2013). Nuclei were extracted from 50,000 naive, effector or memory, sort-purified OT-1 cells and immediately resuspended in transposition reaction mix (Illumina Nextera DNA Sample Preparation Kit - Cat #FC121- 1030) for 30 minutes at 37C. Transposed DNA was purified using a QIAGEN MinElute PCR Purification kit (Cat #28004), and amplified for 5 PCR cycles using PCR primer 1 (Ad1_noMX) and an indexed PCR primer. Aliquots of each amplicon were used as template in a real-time quantitative PCR for 20 cycles to determine the optimal cycle number for library amplification, with amplicons purified as previously. Library quality was determined using a Bioanalyzer (Agilent) to ensure that amplicons ranged between 50-200bp, and samples were subjected to paired-end sequencing on an Illumina Hiseq2500 instrument. Sequence data was mapped to UCSC mm10, then filtered for PCR duplicates and blacklisted regions, then shifted using Alignment Sieve (deepTools; (Ramirez et al., 2014)) and lastly peaks were called with MACS2 (https://github.com/macs3-project/MACS).

### ChIP and FAIRE

Effector T cells were crosslinked with 0.6% formaldehyde for 10 min at RT. Following sonication, immune-precipitation was performed with anti-H3K4me3, H3K27me3 or HK3K27Ac ChIP-grade antibodies and Protein A magnetic beads (Millipore). FAIRE was performed on samples fixed and sonicated as per ChIP, with accessible chromatin extracted twice with phenol:chloroform:isoamyl (25:24:1) (Sigma). FAIRE enrichment was normalised against a total input for which reverse cross-linking had been performed. ChIP and FAIRE enrichment was measured using quantitative real-time PCR, with data normalised against a total input and no-antibody control. Primers used in these assays were reported previously (Russ et al., 2017).

### Hi-C

Hi-C was performed as per Rao (Rao et al., 2014), with the following adjustments: Step 2 - cells were fixed with 1.5% formaldehyde for 10 min at RT. Step 7 - the nuclei extraction buffer contained 0.4% Igepal. Step 12 - restriction digestion was performed overnight with 400U Mbo1 (NEB) in NEB buffer 2.1. Steps 28-35 were skipped. Step 54 - 2.5ul NEBNext Adapter for Illumina (cat # E7370) was used in place of Illumina indexed adapter, with ligation at 20C for 15 min, followed by addition of 3U USER enzyme and further incubation at 37C for 15min. Step 60 - samples were incubated at 95C for 10 min in a thermocycler, and beads were removed before final library amplification with NEBNext Multiplex Oligos for Illumina (cat # E7335S).

### Data normalisation, Differential Loop calling, Gene assignment, MDS plots, GSEA

Raw Hi-C FASTQ files were aligned to the mouse genome (mm10 build), and binned Hi-C matrices generated using Juicer (Durand et al., 2016). Hi-C data was normalised and differential loops were called using multiHiCcompare (Stansfield et al., 2019). TADs were compared using TADCompare (Cresswell and Dozmorov, 2020). GSEA analysis was performed using the FGSEA package (https://www.biorxiv.org/content/10.1101/060012v3), with bubble plots made using a custom Tidyverse script (https://www.tidyverse.org/). MDS plots were generated using the edgeR MDS package (https://rdrr.io/bioc/edgeR/man/plotMDS.DGEList.html). Positioning of dots in MDS is directly proportional to sample similarity.

### Data visualisation

Circos plots were generated using the ShinyCircos package (Yu et al., 2018). Enrichment plots were made using the deepTools2 package (Ramirez et al., 2016). All other figures were made using custom R codes and ggplots2 (Wickham, 2016).

### Flow Cytometry

Single-cell suspensions from spleens, lymph nodes or bronchiolar lavage fluid (BAL) were stained with LIVE/DEAD Fixable Aqua Dead Cell Stain (Thermo Fischer Scientific) for 10 mins at room temperature. Cells were washed with MACS buffer (2mM EDTA, 2% BSA in PBS) prior to resuspension in antibody cocktail containing fluorochrome conjugated antibodies specific for CD4, CD8𝛼, CD45.1, or CD44. For cytokine staining, cells were fix and permeabilised according to manufacturer’s instructions (BD Biosciences) prior to staining with anti-CCL4 and CCL5 antibodies. Stained cells were washed twice with permeabilization buffer, and twice with MACS buffer before analysis. Samples were read with a FACSCanto II cytometers (BD Biosciences), and analysed using FlowJo software (Tree Star, Ashland, OR, USA).

## QUANTIFICATION AND STATISTICAL ANALYSIS

In Figure 2, RNA-seq data are shown as the mean of 2 (memory) or 3 (naive and effector) biological replicate values, ± SEM. In Figure 3A, numbers of *cis* interactions unique to each differentiation state were determined by pairwise comparisons using multiHiCcompare (50Kb resolution, 0.05 FDR). 3B) GSEA analysis comparing genes connected by loops enriched in one condition over another (Y axis), against RNA-seq data derived from matching samples (Russ et al., 2014). Circle sizes reflect adjusted p values (-log10) and colour represents normalised enrichment score (NES), with red indicating enrichment versus the first RNA-seq condition listed in pairwise comparison, and blue indicating enrichment is the second RNA-seq condition listed. In Figure 4 enrichment of transcription factor binding at enhancers unique to naïve or effector (Russ et al., 2017) was performed using curated transcription factor ChIP-Seq data through the CistromeDB Toolkit with shading reflecting GIGGLE score (Zheng et al., 2019). In Figure 5 and 6, GSEA analysis comparing genes connected by loops gained in naïve Bach2^-/-^ and *Satb1^m1Anu/m1Anu^* CD8^+^ T cells, respectively, relative to naïve WT CD8^+^ T cells against RNA-seq data derived from naïve and effector CTLs samples (datasets as described above). p values and normalised enrichment score (NES) are shown (Subramanian et al., 2005). All other Methods used to quantify and perform statistical analyses on data are described in figure legends.

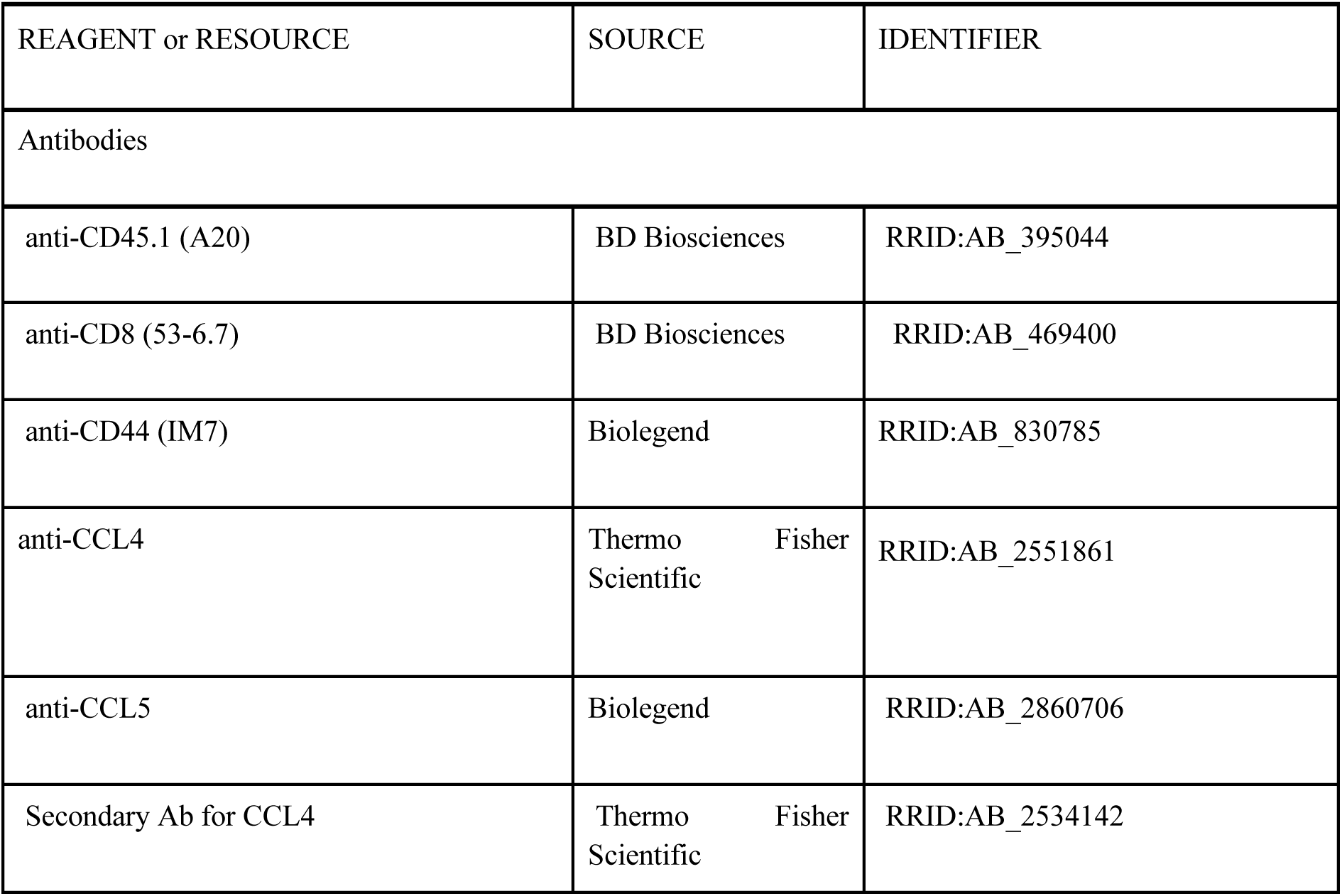
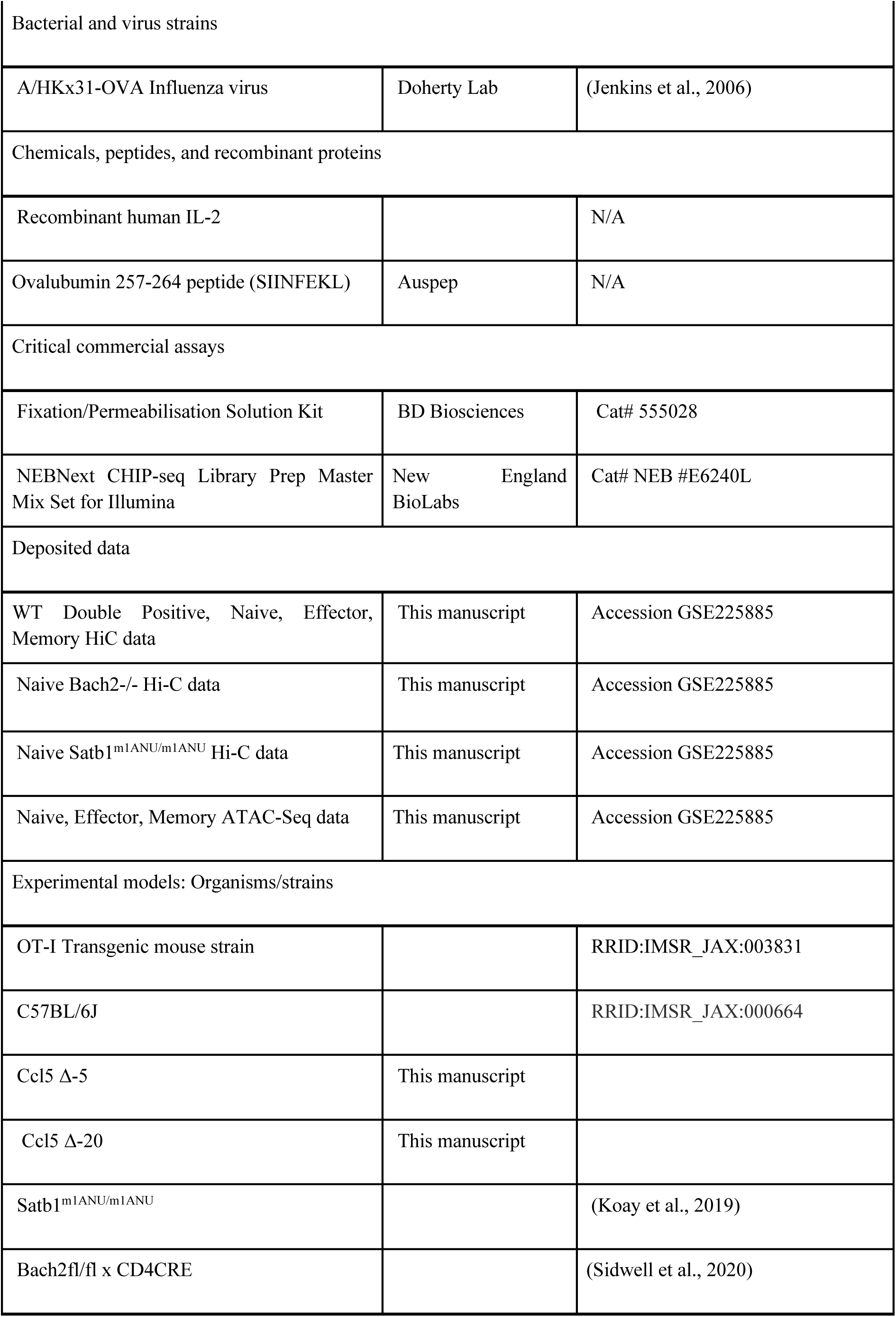
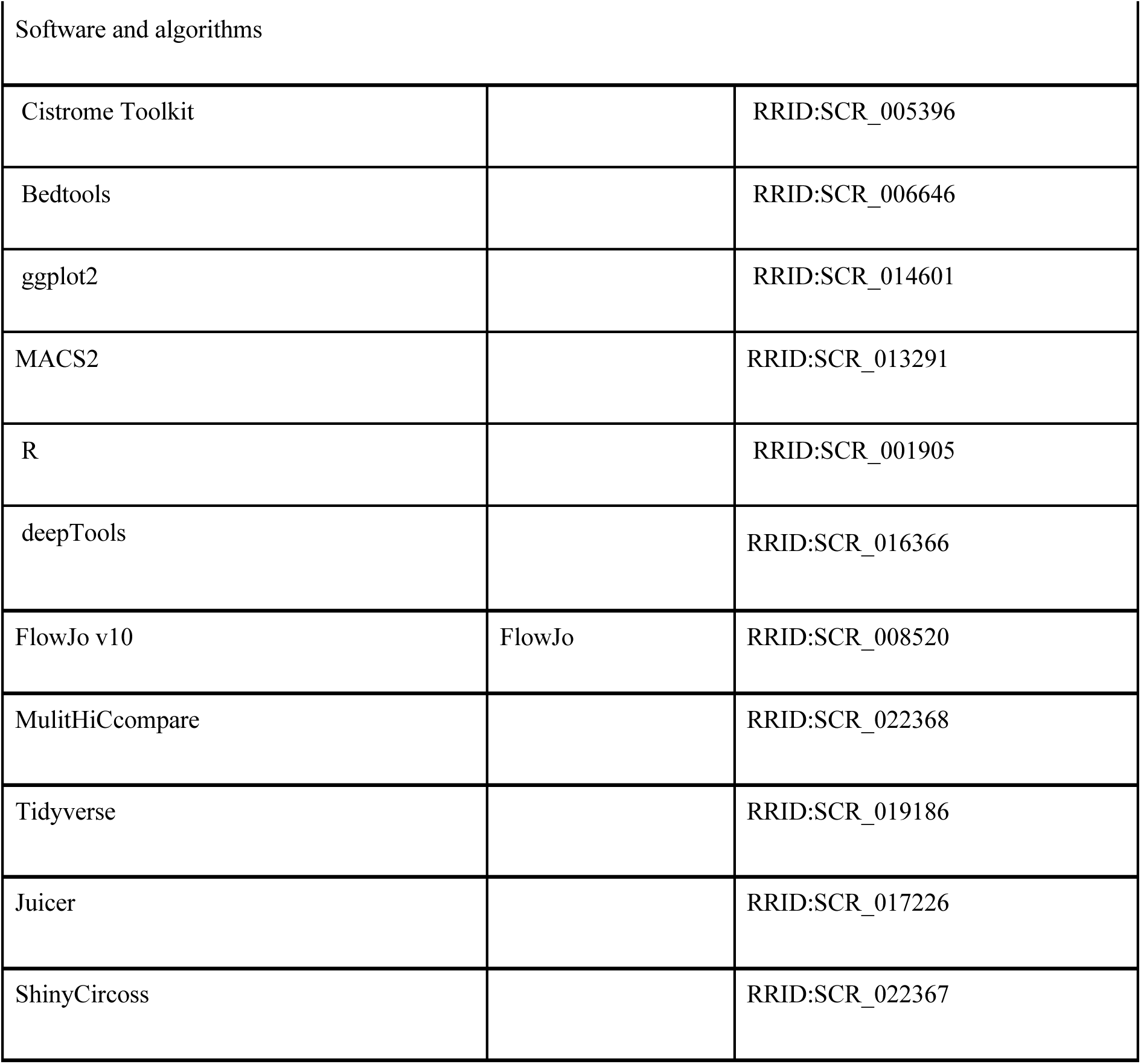

