## Supplemental Figures and Information for "Active maintenance of CD8^+^ T cell naïvety through regulation of global genome architecture"

| REAGENT or RESOURCE | SOURCE | IDENTIFIER |
|---|---|---|
| Antibodies | | |
| anti-CD45.1 (A20) | BD Biosciences | RRID:AB_395044 |
| anti-CD8 (53-6.7) | BD Biosciences | RRID:AB_469400 |
| anti-CD44 (IM7) | Biolegend | RRID:AB_830785 |
| anti-CCL4 | Thermo Fisher Scientific | RRID:AB_2551861 |
| anti-CCL5 | Biolegend | RRID:AB_2860706 |
| Secondary Ab for CCL4 | Thermo Fisher Scientific | RRID:AB_2534142 |

| | | |
|---|---|---|
| Bacterial and virus strains | | |
| A/HKx31-OVA Influenza virus | Doherty Lab | (Jenkins et al., 2006) |
| Chemicals, peptides, and recombinant proteins | | |
| Recombinant human IL-2 | | N/A |
| Ovalubumin 257-264 peptide (SIINFEKL) | Auspep | N/A |
| Critical commercial assays | | |
| Fixation/Permeabilisation Solution Kit | BD Biosciences | Cat# 555028 |
| NEBNext CHIP-seq Library Prep Master Mix Set for Illumina | New England BioLabs | Cat# NEB #E6240L |
| Deposited data | | |
| WT Double Positive, Naive, Effector, Memory HiC data | This manuscript | Accession xxx |
| Naive Bach2-/- Hi-C data | This manuscript | Accession xxx |
| Naive Satb1$^{m1ANU/m1ANU}$ Hi-C data | This manuscript | Accession xxx |
| Naive, Effector, Memory ATAC-Seq data | This manuscript | Accession xxx |
| Experimental models: Organisms/strains | | |
| OT-I Transgenic mouse strain | | RRID:IMSR_JAX:003831 |
| C57BL/6J | | RRID:IMSR_JAX:000664 |
| Ccl5 Δ-5 | This manuscript | |
| Ccl5 Δ-20 | This manuscript | |
| Satb1$^{m1ANU/m1ANU}$ | | (Koay et al., 2019) |
| Bach2fl/fl x CD4CRE | | (Sidwell et al., 2020) |

| Software and algorithms | | |
|---|---|---|
| Cistrome Toolkit | | RRID:SCR_005396 |
| Bedtools | | RRID:SCR_006646 |
| ggplot2 | | RRID:SCR_014601 |
| MACS2 | | RRID:SCR_013291 |
| R | | RRID:SCR_001905 |
| deepTools | | RRID:SCR_016366 |
| FlowJo v10 | FlowJo | RRID:SCR_008520 |
| MulitHiCcompare | | RRID:SCR_022368 |
| Tidyverse | | RRID:SCR_019186 |
| Juicer | | RRID:SCR_017226 |
| ShinyCircoss | | RRID:SCR_022367 |

Buenrostro, J.D., Giresi, P.G., Zaba, L.C., Chang, H.Y., and Greenleaf, W.J. (2013). Transposition of native chromatin for fast and sensitive epigenomic profiling of open chromatin, DNA-binding proteins and nucleosome position. Nat Methods *10*, 1213-1218.
Cresswell, K.G., and Dozmorov, M.G. (2020). TADCompare: An R Package for Differential and Temporal Analysis of Topologically Associated Domains. Front Genet *11*, 158.
Durand, N.C., Shamim, M.S., Machol, I., Rao, S.S., Huntley, M.H., Lander, E.S., and Aiden, E.L. (2016). Juicer Provides a One-Click System for Analyzing Loop-Resolution Hi-C Experiments. Cell Syst *3*, 95-98.
Jenkins, M.R., Webby, R., Doherty, P.C., and Turner, S.J. (2006). Addition of a prominent epitope affects influenza A virus-specific CD8+ T cell immunodominance hierarchies when antigen is limiting. J Immunol *177*, 2917-2925.
Koay, H.F., Su, S., Amann-Zalcenstein, D., Daley, S.R., Comerford, I., Miosge, L., Whyte, C.E., Konstantinov, I.E., d'Udekem, Y., Baldwin, T., *et al.* (2019). A divergent transcriptional landscape underpins the development and functional branching of MAIT cells. Sci Immunol *4*.
Ramirez, F., Dundar, F., Diehl, S., Gruning, B.A., and Manke, T. (2014). deepTools: a flexible platform for exploring deep-sequencing data. Nucleic Acids Res *42*, W187-191.

Ramirez, F., Ryan, D.P., Gruning, B., Bhardwaj, V., Kilpert, F., Richter, A.S., Heyne, S., Dundar, F., and Manke, T. (2016). deepTools2: a next generation web server for deep-sequencing data analysis. Nucleic Acids Res *44*, W160-165.
Rao, S.S., Huntley, M.H., Durand, N.C., Stamenova, E.K., Bochkov, I.D., Robinson, J.T., Sanborn, A.L., Machol, I., Omer, A.D., Lander, E.S., and Aiden, E.L. (2014). A 3D map of the human genome at kilobase resolution reveals principles of chromatin looping. Cell *159*, 1665-1680.
Russ, B.E., Olshanksy, M., Smallwood, H.S., Li, J., Denton, A.E., Prier, J.E., Stock, A.T., Croom, H.A., Cullen, J.G., Nguyen, M.L., *et al.* (2014). Distinct Epigenetic Signatures Delineate Transcriptional Programs during Virus-Specific CD8(+) T Cell Differentiation. Immunity *41*, 853-865.
Russ, B.E., Olshansky, M., Li, J., Nguyen, M.L.T., Gearing, L.J., Nguyen, T.H.O., Olson, M.R., McQuilton, H.A., Nussing, S., Khoury, G., *et al.* (2017). Regulation of H3K4me3 at Transcriptional Enhancers Characterizes Acquisition of Virus-Specific CD8(+) T Cell-Lineage-Specific Function. Cell Rep *21*, 3624-3636.
Sidwell, T., Liao, Y., Garnham, A.L., Vasanthakumar, A., Gloury, R., Blume, J., Teh, P.P., Chisanga, D., Thelemann, C., de Labastida Rivera, F., *et al.* (2020). Attenuation of TCR-induced transcription by Bach2 controls regulatory T cell differentiation and homeostasis. Nat Commun *11*, 252.
Stansfield, J.C., Cresswell, K.G., and Dozmorov, M.G. (2019). multiHiCcompare: joint normalization and comparative analysis of complex Hi-C experiments. Bioinformatics *35*, 2916-2923.
Subramanian, A., Tamayo, P., Mootha, V.K., Mukherjee, S., Ebert, B.L., Gillette, M.A., Paulovich, A., Pomeroy, S.L., Golub, T.R., Lander, E.S., and Mesirov, J.P. (2005). Gene set enrichment analysis: a knowledge-based approach for interpreting genome-wide expression profiles. Proc Natl Acad Sci U S A *102*, 15545-15550.
Wickham, H. (2016). ggplot2 : Elegant Graphics for Data Analysis. In Use R!, (Cham, Springer International Publishing : Imprint: Springer,), pp. 1 online resource (XVI, 260 pages 232 illustrations, 140 illustrations in color.
Yu, Y., Ouyang, Y., and Yao, W. (2018). shinyCircos: an R/Shiny application for interactive creation of Circos plot. Bioinformatics *34*, 1229-1231.
Zheng, R., Wan, C., Mei, S., Qin, Q., Wu, Q., Sun, H., Chen, C.H., Brown, M., Zhang, X., Meyer, C.A., and Liu, X.S. (2019). Cistrome Data Browser: expanded datasets and new tools for gene regulatory analysis. Nucleic Acids Res *47*, D729-D735.

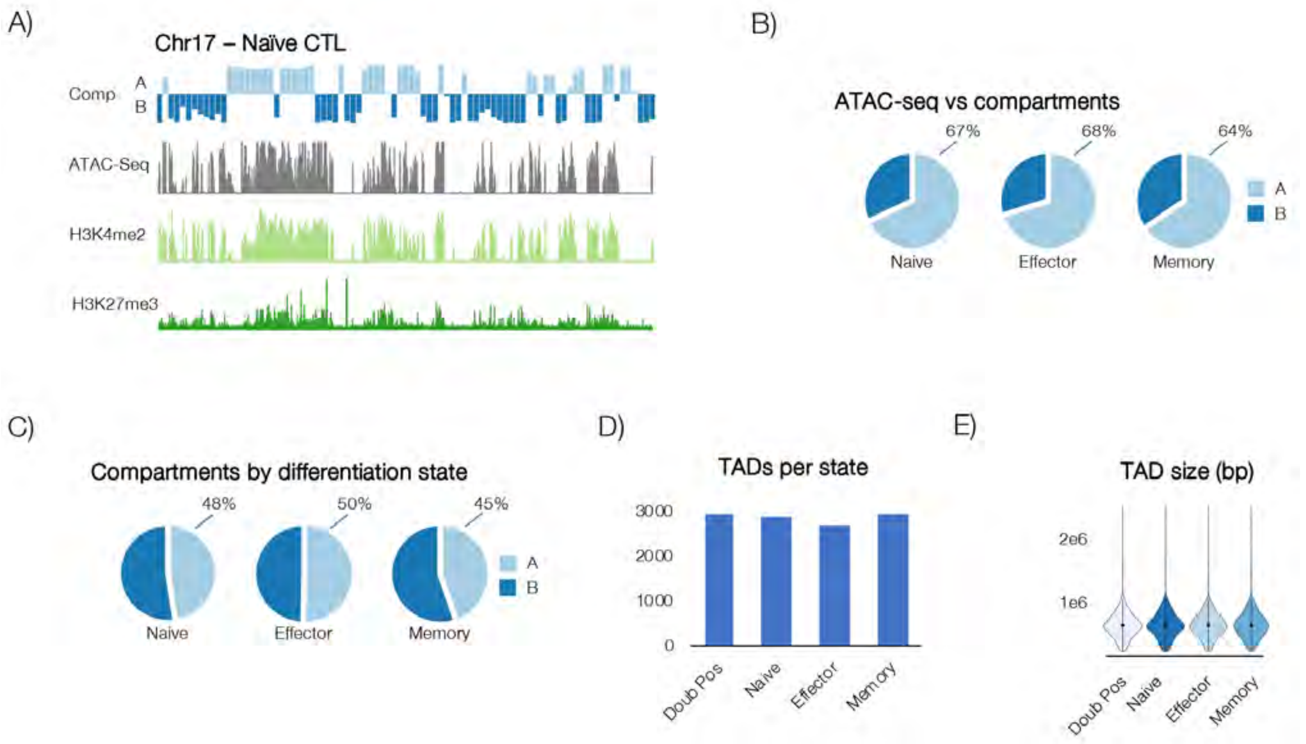


**Supplementary Figure 1. Conservation of higher order chromatin structures during CTL differentiation.** A) Genomic compartments broadly reflect chromatin state. Eigenvectors calculated at 1Mb resolution for chromosome 17 of naïve CTL, with A and B compartments shown in light blue and dark blue, respectively, compared with paired ATAC-seq (grey), H3K4me2 (light green) and H3K27me3 (dark green) ChIP-Seq data. B) Genome-wide overlap of open chromatin peaks (identified by ATAC-Seq) and A/B compartments shows that most open regions occur within the A compartment (indicated by percentage figures). C) Similar A/B compartment distribution between differentiation states. D) Average TAD number and size (E) was stable during differentiation of double positive to naïve and virus-specific effector and memory CTL.

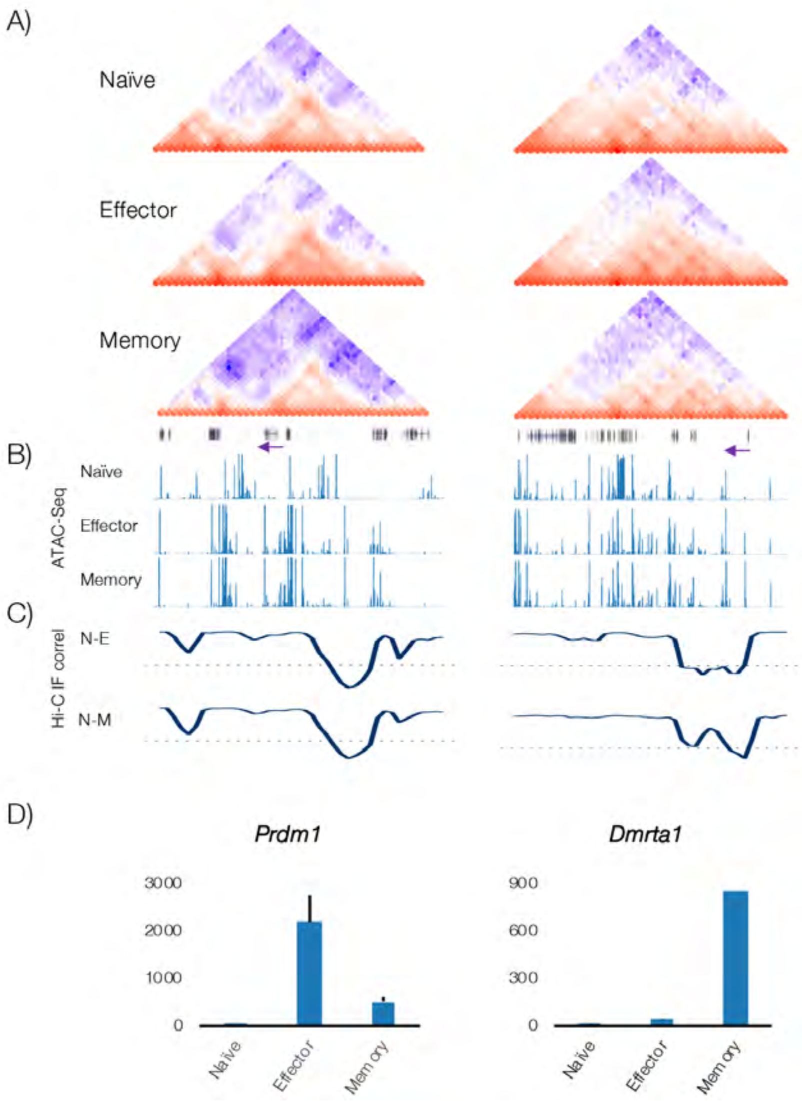


**Supplementary Figure 2.** A) Hi-C data (50Kb bins) normalised using ICED method showing interaction frequency at *Prdm1* and *Dmrta1* loci in naïve, effector and memory OT-1 CTLs. Track below memory panel shows genes, with purple arrow highlighting genes of interest and their direction of transcription. B) ATAC-Seq data for Naïve, effector and memory OT-1 CTLs. C) Pairwise correlation of binned interaction frequencies (50kb) for naïve and effector (N-E), and naïve and memory (N-M) samples, with dotted line indicating 0 on the y axis. D) Bottom panel shows normalised RNA-Seq counts (Russ *et al*, 2014).

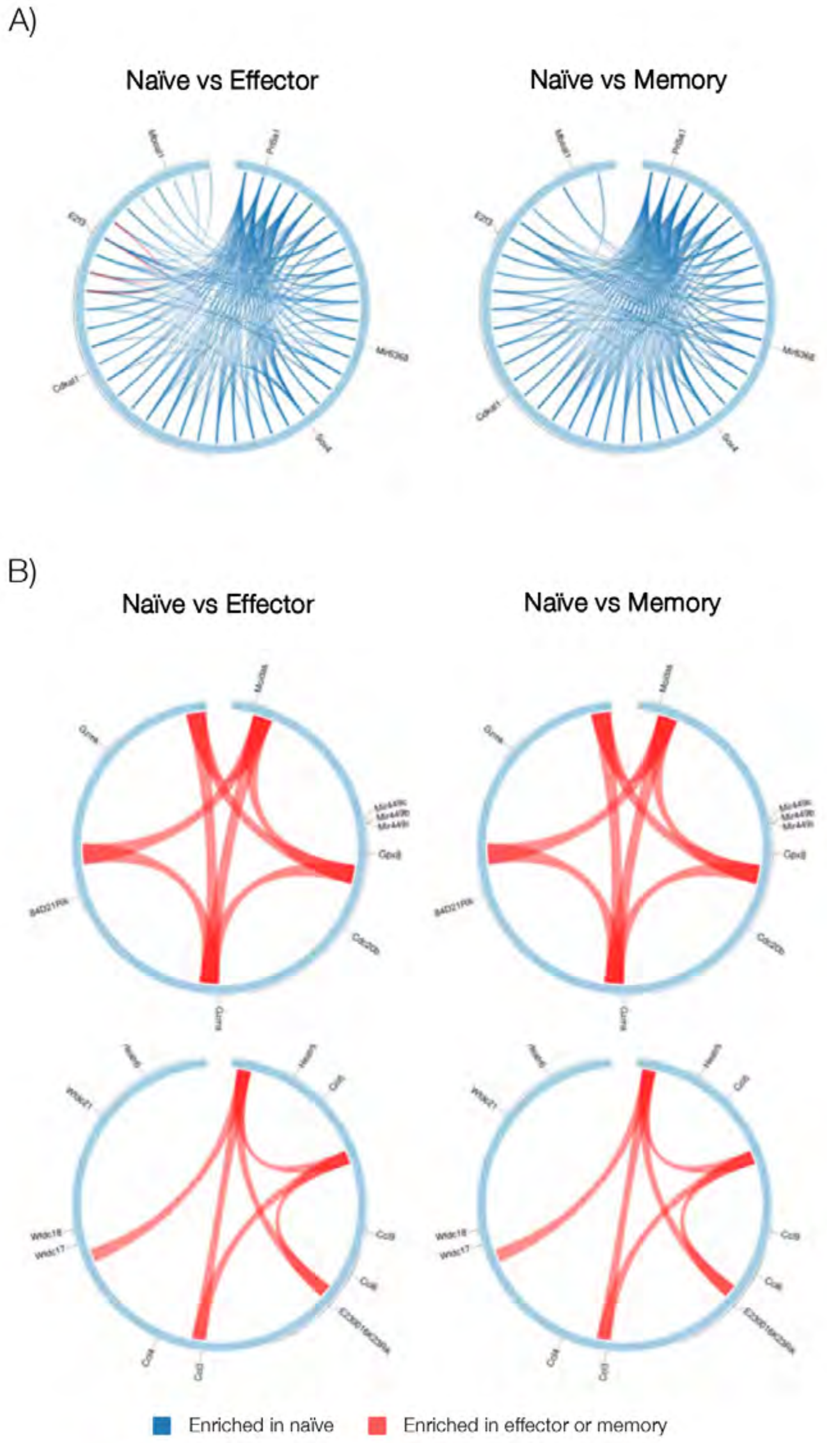


**Supplementary Figure 3. Loss and gain of *cis* regulatory interactions underscores CTL differentiation state specific gene transcription profiles.** A) Examples of loci where loops were largely lost upon differentiation (blue loops are present in naïve over effector or memory; red loops are gained on differentiation). B) Examples of loci where loops were largely gained upon differentiation.

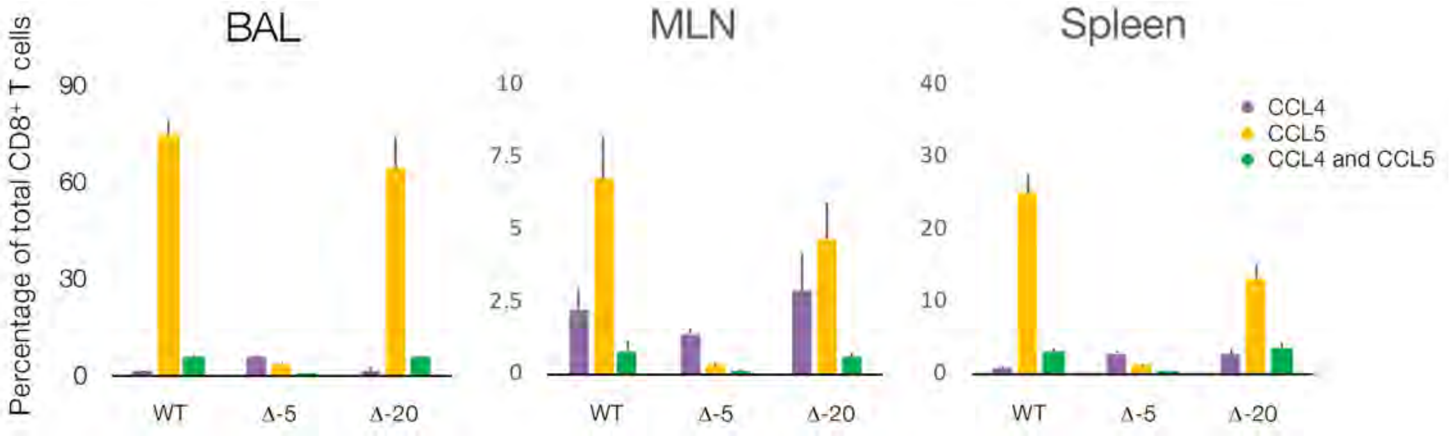


**Supplementary Figure 4. Altered T cell chemokine expression in mice following deletion of cis interacting elements mapped by Hi-C.** Percentage of $CD8^+$ and $CD4^+$ T cells expressing chemokines CCL4 and CCL5 in BAL, Spleen and MLN 10 days post challenge. WT (blue), -5 (red), -20 (green). Error bars are SEM. N=18-27 mice, across 3 separate experiments. % - p=<0.05 WT vs Δ-5; # - p=<0.05 WT vs Δ-20.
